# Enhancing Microalgal Growth through Integrated Computational Modeling and 3D Bioprinting

**DOI:** 10.64898/2025.12.11.693620

**Authors:** Swathi Murthy, Maria Mosshammer, Michael Kühl

## Abstract

Efficient cultivation of microalgae for biofuel and bioproduct applications is often limited by suboptimal light distribution, poor mass transfer, and large land footprint requirements in conventional flat biofilm and open-pond systems. In this study, we combine computational modeling, 3D bioprinting, and experimental measurements to design and evaluate the role of different geometries for photosynthetic production in bioprinted microalgal constructs. Specifically, we investigate a perforated slab with channels (PS) and Vgroove structures, which show enhanced algal growth performance compared to flat slabs of the same volume and footprint. By integrating experimentally measured photosynthetic parameters into radiative transfer and oxygen diffusion–reaction simulations, we uncover the role of structure induced improvements in light penetration, surface area-to-volume ratio, and mass transfer dynamics driving increased algal growth and photosynthesis. Algal growth rates in printed PS and Vgroove geometries were 2.4 and 1.3 times higher, respectively, than in corresponding slab. These findings demonstrate the potential of engineered 3D bioprinted architectures to optimize photosynthetic efficiency and growth rates while minimizing spatial footprint, paving the way for compact, high-throughput bioreactors for sustainable algal cultivation. Beyond biofuel production, these microalgal systems enable sustainable CO₂ capture and utilization.

## 1. Introduction

To meet the world’s growing energy demands sustainably, microalgae have emerged as highly appealing biomass feedstock for biofuel production due to their environmental compatibility, economic viability, renewability, and their ability to produce valuable byproducts.^2–4^ Microalgae can significantly contribute to mitigating global carbon emissions by sequestering CO_2_ during photosynthesis, and they require considerably smaller land areas for biomass production compared to terrestrial plants, primarily due to their superior photosynthetic efficiency, which can be 10 to 50 times higher.^2–4^ Studies highlight that the efficiency of biofuel production from microalgae is strongly dependent on lipid content and biomass productivity, which can reach up to 80% and 7.3 g L^-1^day^−1^ based on the biomass dry weight, respectively, making microalgae ideal candidates for sustainable biofuel feedstock.^2^ Despite these advantages, current algal biofuel production faces economic hurdles due to low biomass productivity, largely resulting from limited utilization of incoming light energy and insufficient mass transfer of nutrients and gasses.^2–4,6^

The choice of cultivation system, such as open ponds or photobioreactors, significantly affects biomass productivity by shaping the algal growth environment, including the management and availability of critical factors such as light, CO_2_, dissolved oxygen, and nutrients.^2,4^ Improving the design of cultivation systems to optimize light harvesting and nutrient mass transfer could thus substantially enhance productivity. In this context, both autotrophic and heterotrophic cultivation strategies offer distinct advantages. Autotrophic systems rely on light and CO₂ and have demonstrated relatively lower production costs per biomass unit, particularly in flat-panel photobioreactors, but they are limited by light availability and large spatial footprints.^7^ In contrast, heterotrophic systems use organic carbon sources such as glucose to achieve significantly higher cell densities and volumetric productivities.^8,9^

Natural phototrophic biofilms face inherent limitations in light penetration, which restricts their photosynthetic efficiency.^10^ This bottleneck can potentially be addressed through advanced photobioreactor designs that optimize both light utilization and mass transfer, ensuring efficient penetration and more uniform distributions of light and nutrients within the algal biomass, improving their photosynthetic efficiency and potential for biofuel production.^2–4,6,11,12^ Inspiration for these innovative structures can be drawn from natural biological systems; for instance, reef-building corals and giant clams that both harbor microalgal symbionts in their tissues and exhibit remarkably high photosynthetic efficiencies due to the efficient light management and mass transfer facilitated by their intricate architecture.^13–15^ Furthermore, detailed simulation studies demonstrate that improved optical configurations and innovative designs can significantly enhance microalgal biomass production, potentially reaching productivity levels above 100 g dry biomass m^−2^ day^−1^ under optimal conditions.^6^ For instance, configurations such as V-shaped bioreactors have experimentally demonstrated significant increases in biomass productivity through effective dispersion and trapping of incident light energy.^6^ The economic viability of microalgal biofuel production is closely tied to the biomass productivity achievable through these optimized cultivation systems, which can lower production costs and enhance the overall sustainability of biofuel processes.^4^ Consequently, predictive modeling and simulation of photophysiological performance become essential tools in designing and optimizing platforms for algal production, facilitating enhanced growth dynamics, and ultimately, commercial scalability of biofuels derived from microalgae. ^6,16^

Computational methods and algorithms for simulating light propagation in tissues have been developed in biomedical optics^17–20^ and can be combined with hydrodynamic and mass transfer simulations^21,22^ to facilitate the design of optimized structures for efficient light utilization and mass transfer. These computational designs can be rapidly prototyped using methods such as 3D bioprinting^16,23–25^, a technique widely used in tissue and organ engineering due to its ability to precisely pattern biomaterials containing human and other mammalian cells in hydrogels.^26–29^. In comparison, the application of bioprinting in other areas of science and engineering remain less explored. In addition to optimizing 3D structural designs, exploring algal cell aggregation and the embedding of scattering particles in the surrounding medium can significantly enhance light management, resulting in up to a fourfold improvement in photosynthetic efficiency and biomass production, as compared to traditional biofilm systems^12^. Recent advances in 3D bioprinting combined with non-invasive imaging and computational simulations can now enable precise control and monitoring of microalgal constructs, optimizing designs for improved photosynthetic efficiency and biomass productivity. ^16^

Here, we present a proof-of-concept study combining computational simulations, 3D bioprinting, and algal metabolism measurements to design and validate geometric structures that significantly enhance algal growth rates compared to conventional flat structures typically seen in natural biofilms^30^ and open-pond cultivation systems^2,3^ for biofuel production. The goal is to engineer space-efficient, structured geometries that maximize light capture and mass transfer, unlocking high-density algal growth within a minimal land footprint. The demonstrated fabrication and simulation pipeline can be broadly applied to investigate other structure-function relationships observed in nature^31,32^, such as symbiotic interactions^33^, and microbe-host dynamics^34–36^, facilitating predictive design and a deeper understanding of the underlying biological mechanisms.

## 2. Methods

Gelatin from porcine skin (gel strength 300, Type A), methacryl anhydride (containing 2,000 ppm topanol A as inhibitor, 94%), NaHCO_3_, and poly(1-vinylpyrrolidone-co-styrene) (38% emulsion in H_2_O, <0.5 μm particle size) were purchased from Sigma Aldrich (sigmaaldrich.com) and used without further modification. The tangential flow filtration (TFF) system was generously provided by PALL (pall.com/), and the MinimateTM TFF capsule with a 10K omega membrane was purchased from PALL. TAP medium was prepared in MilliQ water, according to the recipe: https://utex.org/products/tap-medium?variant=30991736897626#recipe. 0.25% solution of Trypsin-EDTA was purchased from Sigma Aldrich (sigmaaldrich.com). Irgacure 2959 was purchased from BASF (IC2959; BASF, cat. no. 029891301PS04) and Lithium phenyl-2,4,6-trimethylbenzoylphosphinate from Sigma Aldrich (LAP; Sigma Aldrich)

### 2.1 GelMA synthesis

Functionalized gelatin methacryloyl (GelMA) based hydrogels were synthesized according to literature^37^. Briefly, 10 g of gelatin was swelled for 30 minutes at room temperature and fully dissolved at 50 °C in a water bath under stirring in 100 mL MilliQ water. 4 g of methacrylic anhydride was added dropwise to the gelatin solution under vigorous stirring. All the subsequent steps were conducted at low light levels or darkness. The solution was stirred for 1h and centrifuged at 3500 rpm for 3 minutes at room temperature. The pellet was discarded and the clear supernatant was diluted with 2 volumes of deionized water heated to 40 °C. This was followed by dialysis at 40 °C for 5 days using a Minimate^TM^ TFF capsule (10 kDa cut-off). The pH was adjusted to pH 7.4 using NaHCO_3_ (1M), and the solution was filter-sterilized using a vacuum filtration unit with a 0.2 µm filter (PES membrane). The GelMA solution was transferred to 50 mL vials with vented screw caps, snap frozen and lyophilized at −55 °C in a freeze dryer (CoolSafe 55-4 Pro, Ninolab) for 7 days. The lyophilized GelMA was stored at –20 °C prior to usage.

### 2.2 Algal culture

A culture of the green alga *Chlorella sorokiniana* UTEX1230 was maintained in culture tubes with TAP medium (see above) under constant shaking and under a defined photon irradiance from white LED’s (10 μmol photons m^−2^ s^−1^; 400-700 nm) in a temperature-regulated culture cabinet (ALGAETRON, Photon System Instruments) at 24⁰C.

### 2.3 Bio-ink preparation

A photo initiator stock solution was prepared by dissolving Lithium phenyl-2,4,6-trimethylbenzoylphosphinate (LAP; Sigma Aldrich, CAS Number <u>85073-19-4</u>) in 10 mL TAP medium at 70⁰C under constant stirring, where after the solution was allowed to cool to room temperature.^37^ 8% (w/v) GelMA ink was prepared, by soaking GelMA foam in TAP medium with added photo-initiator stock solution (to final concentration, w/v of 0.25%), overnight in an refrigerator. This was followed by dissolving the GelMA-containing TAP solution in an incubator at 37 ⁰C and stirring at 300 rpm, until the mixture turned clear.^37^ The pH of the solution was adjusted to pH 8.1 by adding a strong base (1M NaOH). The GelMA ink was finally autoclaved at 80 ⁰C for 15 minutes to ensure its sterility. The ink was cooled to <30 ⁰C, before adding the microalgal cells. The algal cells were spun down from the stock solution at 9000 rpm for 60 s. The supernatant was discarded, and the algal cells were added at a concentration of 19 ± 1.2 x 10^6^ cells mL^−1^ bioink to obtain a relatively low biomass load. At ∼26 °C, the hydrogel exhibited low viscosity, allowing the hydrogel–algal cell mixture to be pipetted up and down several times to achieve homogeneous cell distribution within the bioink.

### 2.4 Bioprinting

Three different construct geometries with similar volume and food print, i.e., a construct with V-grooves^6^ (30⁰ vertex angle, 4.7 mm height, volume 123.28 mm^3^, footprint 52.34 mm^2^), a perforated slab construct (volume 175.64 mm^3^, footprint 72.25 mm^2^) and a corresponding simple slab structure, were designed in CAD software (Autodesk Fusion 360.ink), followed by slicing and g-code creation in PrusaSlicer (PrusaSlicer.ink). The g-code was uploaded and printed on a commercial 3D bioprinter (Bio-X, CellInk Lifesciences), which utilizes an extrusion-based printing technique. The printing was carried out with an infill density set to 60% using a 3D honeycomb pattern. The print speed and extrusion pressure were set to 2 mm s^−1^ and 25 kPa, respectively. The temperature of the bio-ink was set to 24 ⁰C, to obtain a desired viscosity for printing. The constructs were printed, on a PET foil substrate maintained at 10⁰C and were allowed to stand for 5 minutes (after printing the entire construct) to facilitate physical cross-linking, before curing with 405 nm light, at intensity of 3 mW cm^−2^ for 30 seconds. The prints were stored in sterile petri dishes with TAP medium sealed off at the edges with parafilm. The printing and handling were carried out in a sterile environment, either in the bioprinter (fitted with UV-C germicidal lamps and a HEPA H14 dual-filter system) or in a laminar flow bench. The prints were stored in an incubation chamber under constant photon irradiance (PAR; 400-700 nm) of 10 μmol photons m^−2^ s^−1^ from white LED’s (AlgaeTron AG 230) between day 0 and day 1, and under 60 μmol photons m^−2^ s^−1^ between day 1 and day 4.

### 2.5 Biomass quantification via cell counts

The cell density was determined at the beginning of the experiments (day 0) by measuring the cell concentration in the algal stock solution, before adding it to the bio-ink. The cell concentration in the bio-prints on day 4 was determined by removing the prints from the growth medium and dissolving them in 1000 µL trypsin solution^38^ (0.25% Trypsin/EDTA) at 37 ⁰C for 1 hour. For cell counting, the algal solution (stock culture samples or dissolved prints in trypsin) was diluted 3000 to 4000 times in TAP medium, depending on the algal cell concentration. Then 1 mL of the diluted algal solution was poured into a Sedgewick rafter counting chamber, which subdivides 1 mL into (1000) 1µL volume fractions. The number of algal cells in each 1 µL volume fraction was manually counted under an optical microscope. At least 15 to 20 volume fractions were counted for each sample and averaged.

### 2.6 Optical coherence tomography (OCT) measurements and analysis

A spectral domain OCT system (Ganymed II; Thorlabs GmbH, Dachau, Germany) equipped with an objective lens with an effective focal length of 18 mm and a working distance of 7.5 mm (LSM02-BB; Thorlabs GmbH, Dachau, Germany) was used for OCT imaging.^39^ The system is equipped with a 930 nm light source, yielding a maximal axial and lateral resolution in water of 5.8 μm and 8 μm, respectively. Two-dimensional OCT B-scans were acquired at a fixed pixel size of 584 x 1024. The actual field of view was variable in y but fixed in z (= 2.2 mm). The OCT system was optimized to yield highest signal at a fixed distance, in the upper 1/3rd of the image.^39^ OCT imaging was performed on bio-printed constructs fully immersed in TAP medium in a petri-dish. System calibration and optical parameter extraction were performed according to previously published procedures.^16,40–42^

### 2.7 Variable chlorophyll fluorescence imaging

The photosynthetic performance of the microalgae in the bioprinted constructs was measured via a pulse-amplitude-modulated, variable chlorophyll fluorescence imaging system (I-PAM/GFP, Walz GmbH, Effeltrich, Germany) using the pulse-saturation technique.^43,44^ The system utilizes blue (470 nm) LED light for weak (<1 μmol photons m^−2^ s^−1^) modulated measuring light pulses, strong (0.8 s at >2500 μmol photons m^−2^ s^−1^) saturating light pulses, and defined levels of blue actinic irradiance, as measured with a calibrated photon irradiance meter at the level of the bioprinted constructs (ULM, Walz, Effeltrich, Germany).

From measurements of the minimum fluorescence yield, *F_0_*, and the maximum fluorescence yield, *F_m_*, in dark acclimated samples, as recorded before and during a saturation pulse, respectively, we calculated the maximum quantum yield of PSII as *F_v_/F_m_ = (F_m_-F_0_)/F_m_*. This parameter is frequently used to indicate the health and photosynthetic capacity of photosynthetic organisms^45^. The effective PS II quantum yield in light-exposed samples was calculated as *YII = (F^’^ –F)/F^’^*, where *F’* is the maximal fluorescence yield of light acclimated samples (under the saturating light pulse) and *F* is the fluorescence yield under ambient actinic light conditions.^43,46^

From YII measurements after a 10 s exposure at each light level, covering over a range of increasing actinic photon irradiance levels of photosynthetic active radiation (PAR; 400-700 nm), we calculated a so-called rapid light curve (RLC).^46^ The RLC quantifies how the relative electron transport rate (rETR = YII x E_d_) via PSII changes with increasing photon irradiance, E_d_, and can be used as a qualitative measure of the photo-physiological acclimation state of the micro-algae embedded in the prints.^46^

### 2.8 Gas exchange measurements

The net O_2_ exchange between bioprinted constructs and their surrounding medium was measured with samples kept in a customized, gas-tight glass chamber (10 mL volume) with a flat, glass coverslip as cover^47^. The edges of the chamber were painted black, to avoid light guiding and scattering effects. The construct was placed on a small, flat holder inside the chamber above a stirring bar, ensuring constant flow and efficient mass transfer between construct and the surrounding water. An optical O_2_ sensor spot with an optical isolation (PyroScience GmbH) was mounted inside the chamber and could be read out via an optical fiber attached to the chamber at one end and to a fiber-optic O_2_ meter (FireStingO_2_; PyroScience GmbH) at the other end. Calibration of the sensor spots were done according to the manufacturer, and O_2_ concentrations inside the experimental chamber were logged every 10 s enabling a quantification of the net O_2_ exchange between the bioprinted constructs and the surrounding water in the closed chamber.

The net O_2_ production, J, per constructs, where positive values indicate net production and negative values indicate net consumption of O_2_, was calculated from the measured linear change in O_2_ concentration in the chamber over time, dC/dt, corrected for the water volume surrounding the print (V) and the printed construct volume (V_construct_):

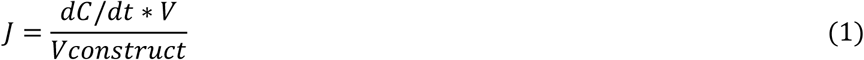

The O_2_ flux was normalized to print volume since the structured print and its corresponding slab had identical volumes, enabling direct comparison of their rates. For recording of O_2_ dynamics, the bioprinted constructs were kept in darkness for 10 minutes prior to switching on the light source, i.e., a fiber-optic tungsten-halogen lamp (K2500 LCD, Schott GmbH). The incident photon irradiance of photosynthetically active radiation (PAR; 400-700 nm) at the level of the construct surface in the chamber was measured with a calibrated spectroradiometer (BTS 256, Gigahertz Optics GmbH) for different lamp settings. The bioprints were then exposed to increasing photon irradiance (100, 200, 270, 430 and 530 µmol photons m^−2^ s^−1^; 400-700 nm) with intermittent dark periods for about 18 minutes each. Measurements were logged continuously every 10 seconds. The flux measurements were used to determine the net production (NP) and dark respiration (DR) rates after a period of illumination, where after the gross photosynthesis rate was estimated as (NP + DR). The different constructs were measured simultaneously in separate vials. The net production of O_2_ as a function of incident photon irradiance was fitted to an exponential function according to Spilling et al.^1^ and the estimated gross photosynthesis according to Webb et al.^5^

### 2.9 Approximation of optical properties for simulation

The optical properties of bio-prints used for simulating the light distribution (see 2.11) were approximated from either the cell count or the OCT data (section 2.6). The algal cell count was used to calculate the absorption coefficient, µ_a_, and the scattering coefficient, µ_s_, based on an earlier experimental studies on the green microalga *Chlorella vulgaris*^48^. To account for the algal growth dynamics, calibrated OCT images (Figure 3) were used to extract the scattering coefficient, µ_s_, and the scattering anisotropy factor, g, for samples on day 4 (summarized in Table S1). It was assumed that the extracted µ_s_ at 930 nm would be similar to the value at 636 nm (used for simulations).

### 2.10 Calculation of net-photosynthesis quantum efficiency

The net-QE for all the samples on day 4 was calculated under an incident photon irradiance of 430 µmol photons m^−2^ s^−1^ (400-700 nm), from the gas exchange data shown in Figure 3b and 3d. At the selected photon irradiance, all samples exhibited net O_2_ production. The following equation was used to calculate the net-QE^49^:

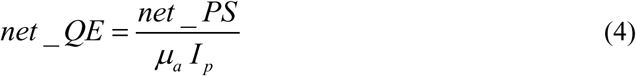

where net_*QE* is the quantum efficiency (net production) of algal photosynthesis, net_PS is net production of O_2_ obtained from respirometery data (figure 3b and 3d), µ_a_ is the absorption coefficient of the bioprint obtained from cell count (figure S7), *I_p_* is the photon scalar irradiance.

### 2.11 Monte Carlo simulation of radiative transfer in bioprinted constructs

Monte Carlo (MC) modelling was used to calculate the radiative transfer in the bioprinted constructs^50^ using the free finite-element MC simulation software called ValoMC^18^, employing the algorithm described by Prahl *et al*.^51^, was used in combination with COMSOL Multiphysics (v5.6, COMSOL Inc., Burlington, MA) as described in our previous study^16^. Briefly, the discretized three-dimensional geometry (created in COMSOL, minimum tetrahedral element size 3 µm) was loaded into ValoMC. A set of material properties was assigned to each of the model domains (summarized in Table S1), consisting of *μ_a_*, *μ_s_*, *g* (using the Henyey-Greenstein approximation for the scattering phase function), and the refractive index *n* (1.33). A ‘direct’ light source at 636 nm, was assigned over the entire top boundary (as shown in figure S2) with a total power of 1 W. For each simulation, 10^8^ photon packets were launched. The output scalar irradiance from the MC simulation was normalized to the was normalized to the incident photon irradiance. The normalized light field was imported into COMSOL and mapped over the 3D bio-print model.

The photon scalar irradiance *I_p_* was calculated from normalized scalar irradiance (*I_s_*), as shown in equation (2).

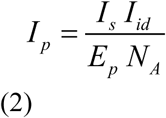

where *I_id_*is the incident downwelling irradiance, *N_A_* the Avogadro’s number, and *E_p_* the energy of a photon at 636 nm, (λ), calculated as *h*c/ λ, *h* is Planck’s constant and *c* is the velocity of light in vacuum.

### 2.12 Mass transfer simulation

Mass transfer simulations were carried out in COMSOL Multiphysics (v5.6, COMSOL Inc., Burlington, MA).

#### Fluid flow

Stationary and incompressible Navier-Stokes equations for laminar flow were used to simulate the water flow over the bio-print in a flow chamber (figure S1), as described in previous work.^16^. A constant water density and dynamic viscosity at 25°C was used. A fully developed laminar flow with average velocity of 5 mm s^−1^ was assigned to the inlet. The transport of the photosynthetically produced O_2_ in the bioprint, was supported by the water velocity profile calculated over the bio-print (described under oxygen transport and reactions). A mesh-independent study (Figure S2) showed that the chosen mesh size, minimum element size of 4 µm for light and 82 µm for O_2_ simulation, is sufficient.

#### Oxygen transport and reactions

The dissolved oxygen concentration (*c_O2_*), was calculated from diffusion-reaction equations in the bio-print and from diffusion-convection equation in the water column, as explained in our previous work^16^. The net rate of O_2_ produced by photosynthesis in the bio-print was calculated by

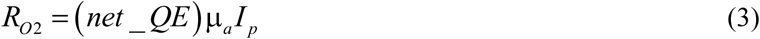

where net_*QE* is the quantum efficiency (net production) of algal photosynthesis (assumed to be 0.005 for day 0 or calculated from respirometery measurements on day 4), µ_a_ is the absorption coefficient of the bioprint, *I_p_* is the photon scalar irradiance.

## 3. Results and discussion

Using predictive modeling we demonstrated how the integration of simulation tools, 3D bioprinting, and algal metabolism measurements can inform the design of geometries that enhance algal growth compared to flat structures. Assuming a specific algal biomass concentration and distribution, used to determine optical properties^48^, we designed two structures: a perforated slab (PS) and a V-groove geometry (Figure 1a). These designs were selected based on their potential to improve light penetration and/or mass transfer relative to a flat slab of the same volume and footprint (Figure S3).

**Figure 1:**
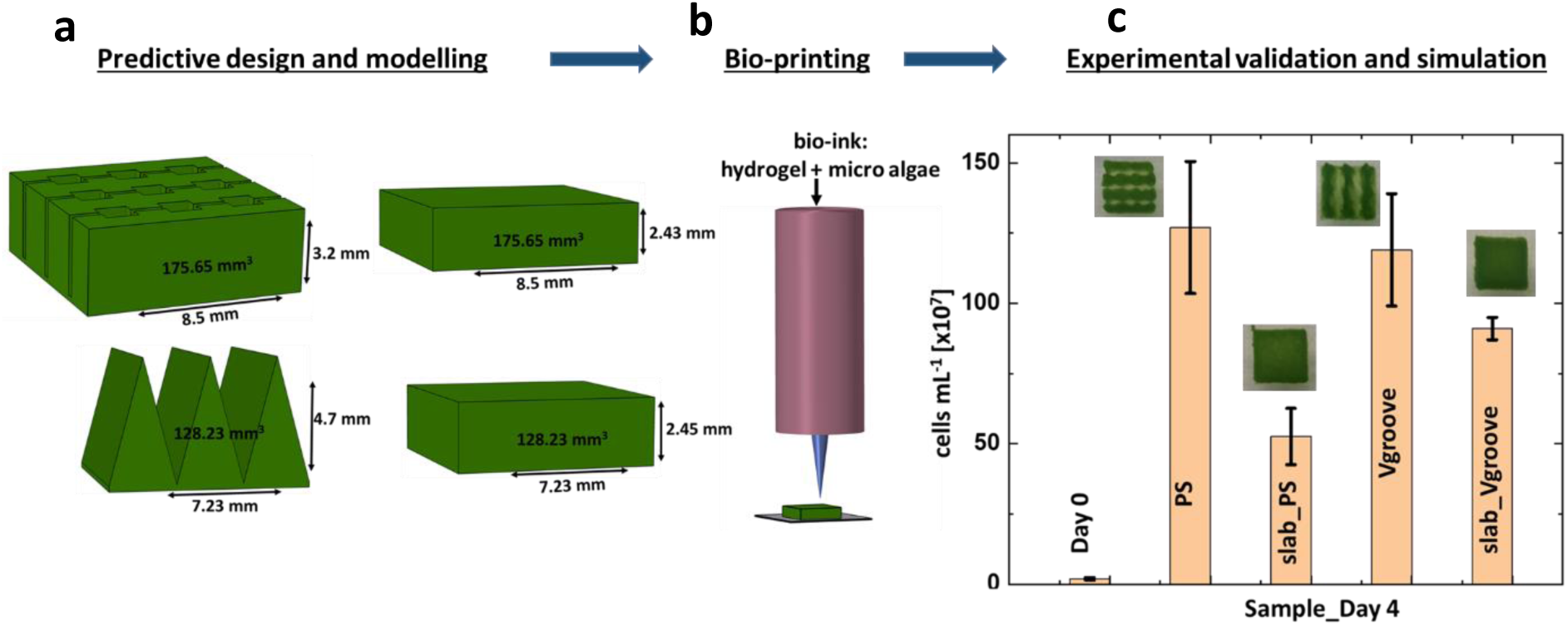
Schematic representation of design-construction-evaluation pipeline. a: Predictive designs of algal bioprint geometry for enhanced algal growth rates comprising a perforated slab (with holes connected by channels), a Vgroove design and the corresponding slabs of same volume and footprint for comparison; b: Bioprinting of the designs in GelMA hydrogel mixed with microalgae; c: Algal cell counts in the different samples on day 4 after printing. The inserts are photographs of the samples (in medium) on day 4.

The corresponding bioprinted samples (Figure 1b, S4) were incubated in growth medium under constant illumination with an incident photon irradiance (400-700 nm) of 60 μmol photons m⁻² s⁻¹ for up to 4 days. Optical and metabolic measurement techniques were used to assess growth performance and extract key parameters for further simulations. Cell count measurements (Figure 1c) confirmed that both structured geometries supported higher algal growth rates than their flat slab counterparts.

Post-experiment simulations, incorporating the measured properties of the printed samples, validated these observations and provided insight into the mechanisms driving enhanced performance. Both light distribution and mass transfer were found to independently contribute to improved growth. The shape of the bioprinted structures influenced the spatial arrangement of algal clusters by modulating local light environments (Figure 3, S5), which in turn affected photosynthetic efficiency.

By day 4, the PS and Vgrove constructs supported an average of 2.4 and 1.3 times more algal biomass, respectively, as compared to their respective flat slabs (Figure 1c). Analysis of the light and mass transfer simulations, incorporating experimental data (Figure S6), suggests that the enhanced growth in PS is primarily driven by high light penetration (comparable to that of the slab on Day 0), along with improved mass transfer and a 2.5-fold increase in exposed surface area relative to the slab. This enhancement likely supports sustained autotrophic growth by maintaining high light penetration. Additionally, the increased surface area and mass transfer could facilitate supplementary heterotrophic metabolism, which has been shown to enable higher cell densities and growth rates under conditions where light becomes limiting^8,9^. Thus, the PS design may leverage both autotrophic and heterotrophic contributions to promote elevated biomass accumulation. The growth advantage in the V-groove design appears to stem from a balance of improved light distribution and more efficient mass transfer as biomass accumulates, supporting both autotrophic activity and complementary heterotrophic metabolism.

### 3.1 Variable chlorophyll fluorescence imaging

Variable chlorophyll fluorescence imaging (VCFI) enables non-invasive monitoring of algal photophysiology with minimal sample manipulation.^16,43,44,46^ Here were used VCFI on Day 1 and Day 4 after bioprinting to monitor microalgal health in the bioprinted constructs and estimate saturating light levels for their photosynthetic activity.

Figures 2a and 2c show the effective quantum yield of photosystem II (YII) and the derived relative electron transport rate (rETR), respectively, plotted against photon irradiance (400-700 nm; μmol photons m⁻² s⁻¹) for all samples on Day 1 and Day 4. Figure 2b presents the maximum quantum yield (F_v_/F_m_) for all samples on both days, with corresponding images shown in Figure S9.

**Figure 2:**
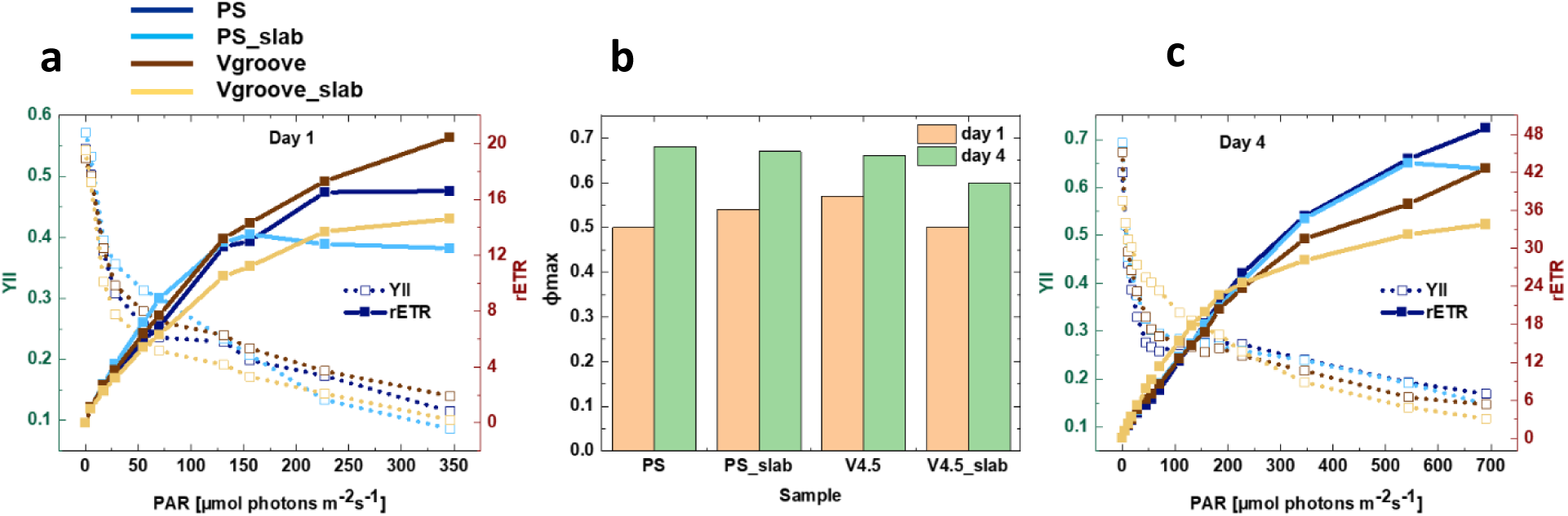
Variable fluorescence imaging of algal bioprints. a: Plot of YII and relative photosynthetic electron transport rate, as integrated over the entire sample surface, versus photon irradiance (μmol photons m^−2^ s^−1^) on day 1. b: Plot of maximum quantum yield (F_v_/F_m_) for dark acclimated samples on day 1 and day 4. c: Plot ofeffective PSII quantum yield (YII) and relative photosynthetic electron transport rate (rETR), as integrated over the entire sample surface, versus photon irradiance (μmol photons m^−2^ s^−1^) on day 4.

**Figure 3:**
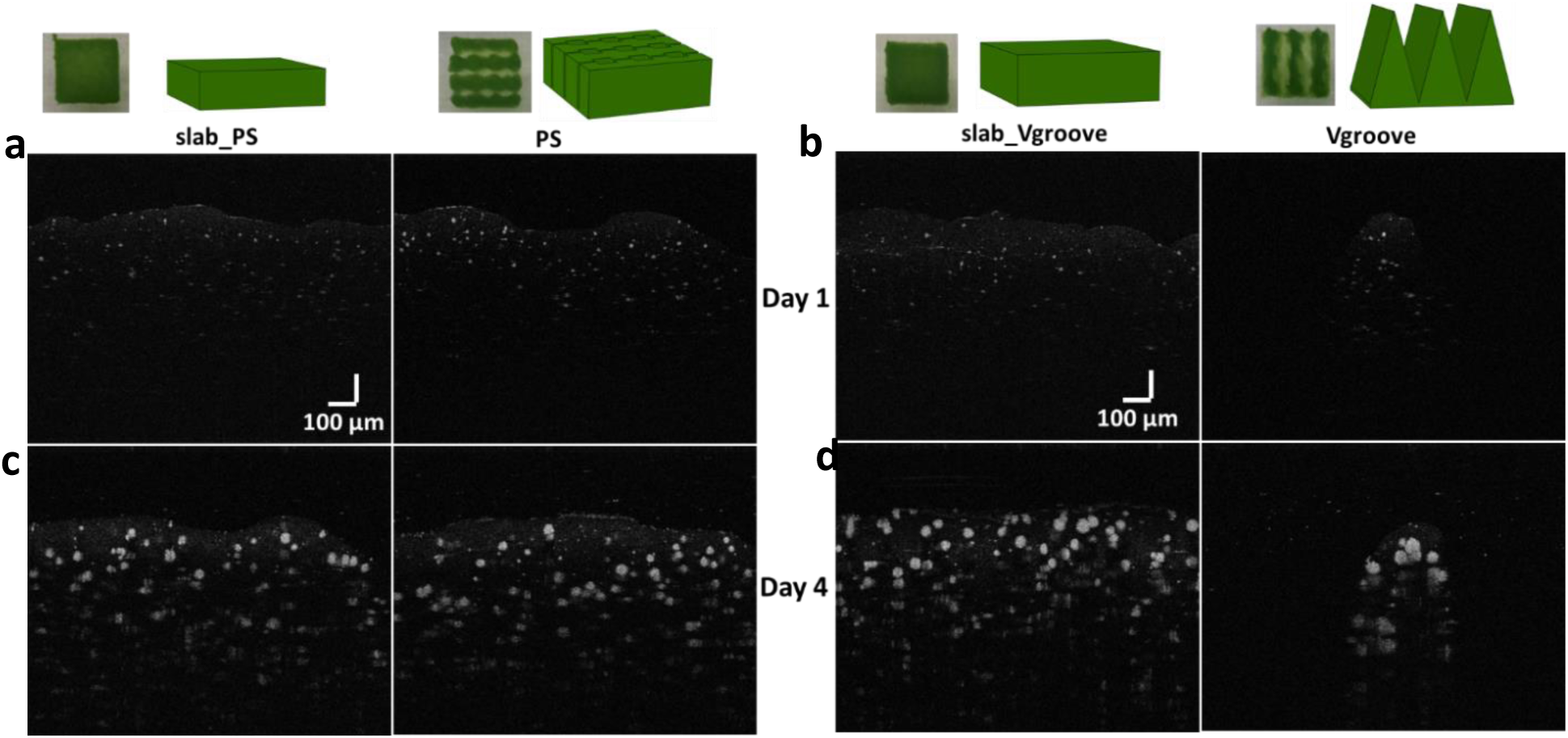
Optical coherence tomography (OCT) images of algal biomass in bioprinted constructs. **a:** slab_PS and PS on Day 1; **b:** slab_Vgroove and Vgroove on Day 1; **c:** slab_PS and PS on Day 4; **d:** slab_Vgroove and Vgroove on Day 4. Inserts on top show 3D sketches and orthogonal images of the corresponding samples.

On Day 1, all samples exhibited a maximum PSII quantum yield, F_v_/F_m_, of approximately 0.5 – 0.55, which increased to 0.6 – 0.7 by Day 4 indicative of increasing photosynthetic capacity following bioprinting. The onset of photosynthetic saturation occurred at around 125 μmol photons m⁻² s⁻¹ on Day 1 and shifted to approximately 200 μmol photons m⁻² s⁻¹ on Day 4. The saturation point is estimated as the inflection or plateau of the YII vs. irradiance curve. These PAM measurements indicate that the microalgae remained physiologically healthy throughout the observation period. The upward shift in saturation light intensity, on Day 4 compared to Day 1, indicates acclimation of the algal photosynthetic apparatus after bioprinting, likely through recovery of PSII function and enhanced light utilization capacity, enabling the cells to tolerate and exploit higher irradiance levels^52^.

### 3.2 Optical Coherence Tomography imaging and extraction of optical properties

Optical Coherence Tomography (OCT) is a non-invasive interferometric imaging technique that captures backscattered photons from internal microstructures and interfaces with refractive index (n) mismatches.^53^ OCT images inherently contain information about the scattering properties of the sample. Using a theoretical model based on the inverse Monte Carlo method, the depth-resolved OCT signal and local signal intensity were fitted to estimate the scattering coefficient (μ_s_) and scattering anisotropy (g).^54^ We imaged the distribution of microalgae within the bioprints using OCT on Day 1 (Figures 3a and 3b) and Day 4 (Figures 3c and 3d). Optical properties were extracted as previously described^16^ (see section 2.9). The resulting parameters are summarized in Figure S1.

As shown in Figure 3 (and the 3D renderings in Figure S5), the algal clusters on Day 1 were relatively small and sparsely distributed. By Day 4, the clusters had grown significantly, forming denser and more highly scattering biomass aggregations (Figure S5). Such growth dynamics affected the internal light distribution within the bioprints, which in turn influences algal growth rates. The extracted optical properties were used in light transport simulations, which serves as inputs for O₂ mass transfer modeling (see Sections 3.4, 2.11, and 2.12).

### 3.3 Respirometry measurements

The light dependent oxygen dynamics within the algal bioprints were measured by determining changes in dissolved oxygen concentration over time in the bulk medium surrounding the printed constructs during incubation in a custom-built gas-exchange chamber^47^ (see section 2.8). As seen in figure 4a and 4c, the structured samples, PS and Vgroove, show a steeper response (net oxygen production rate) with increasing photon irradiance as compared to the respective slabs. The gross photosynthesis was calculated form the measured net photosynthesis rates and the corresponding post-illumination dark respiration rates for each photon irradiance level (Figure S10). The print volume normalized net production rate is plotted in figure 4b and 4d for PS and Vgroove, respectively, along with the respective slabs. All samples exhibit a positive net production of O_2_ above a photon irradiance of 300 µmol photons m^-2^s^−1^. Hence, the measured data point at 430 µmol photons m^-2^s^−1^ was used to calculate the net-quantum efficiency (net-QE) for net photosynthetic O_2_ production, as described in section 2.10. The calculated net-QE for all samples, on Day 4, are summarized in Table T2. This was used to simulate O_2_ mass transfer as described in section 2.12.

**Figure 4:**
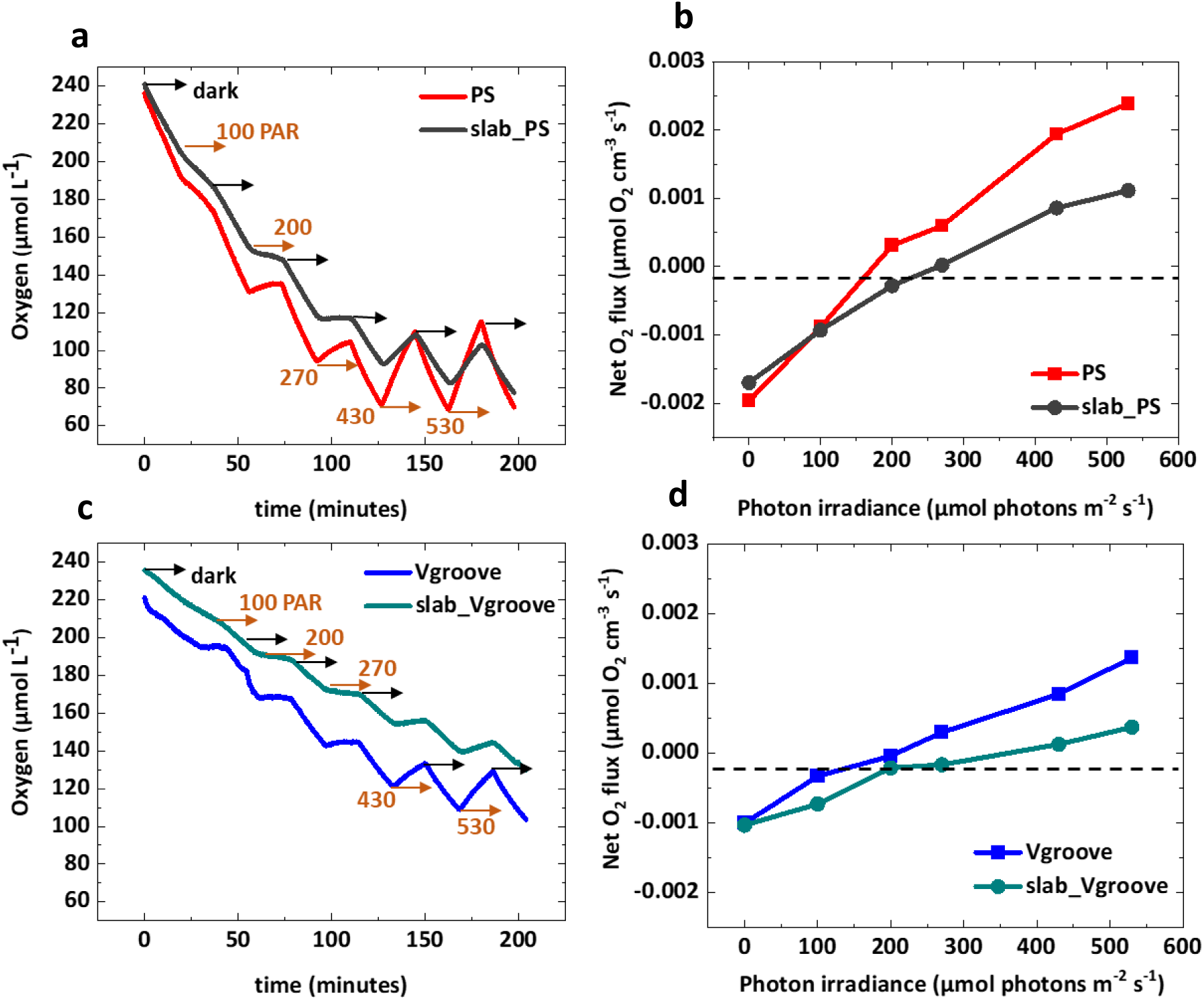
Respirometry measurements of net O_2_ exchange under different light levels (PAR) with alternating dark measurements of. **a:** PS and slab_PS; **c:** Vgroove and slab_Vgroove. Plots of net O_2_ production under different light levels, normalized to print volume, as calculated from linear slopes in a and c for **b:** PS and PS_slab; **d:** Vgroove and slab_Vgroove.

### 3.4 Light and mass transfer simulation

#### 3.4.1 PS versus slab_PS

The simulated steady-state scalar irradiance, when combined with photosynthetic oxygen production and distribution, can help reveal the underlying mechanisms behind the experimental observations. For Day 0, the 2D cross-section of the simulated light field in the constructs (Figure 5a, figure S11a) and the corresponding 1D profiles extracted along a vertical line (Figure 5c, Figure S11c) show that light penetrated through the entire sample. The overall average light intensity (Figure S6a) on day 0 was similar for PS (95% of the slab) and slab_PS. However, the simulations showed a higher surface area of illumination for PS (2.5 times) compared to slab_PS. The 2D cut-plane data of the simulated steady-state O₂ distribution for day 0 (Figure 5b, S11b) and the corresponding 1D profiles (Figure 5c, S11c) indicate higher O₂ accumulation in slab_PS compared to PS. This difference can be attributed to PS having more efficient mass transfer with the surrounding medium, due to its surface area-to-volume ratio being 2.5 times greater than that of slab_PS. As a result, the overall average steady-state O₂ concentration for PS on day 0 is approximately 65% of that observed in slab_PS (Figure S6a).

**Figure 5:**
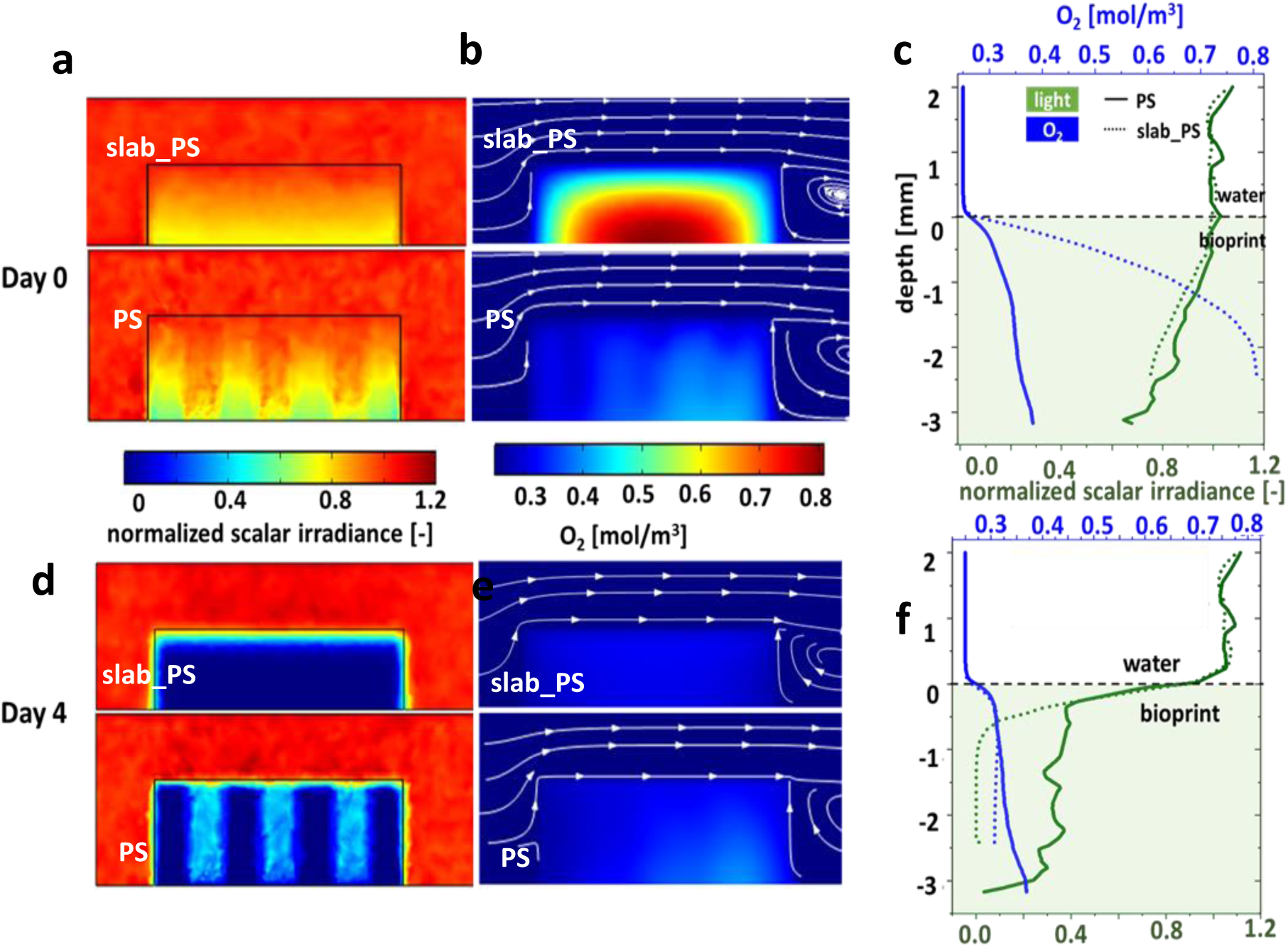
Simulation of steady state light and O_2_ distribution in bioprinted PS and slab_PS constructs. **a:** 2D cross-section (see figure S7) showing the simulated vertical light distribution (normalized scalar irradiance 636 nm) on Day 0; **b:** 2D cross-section of the corresponding O_2_ distribution on Day 0; **c:** Extracted line profiles (see figure S7) of normalized scalar irradiance (relative to incident irradiance) and O_2_ concentration on Day 0. **d:** 2D cross-section (see figure S7) showing the simulated vertical light distribution (normalized scalar irradiance for 636 nm) on Day 4; **e:** 2D cross-section of the corresponding O_2_ distribution on Day 4. **f:** Extracted line profiles (see figure S7) of normalized scalar irradiance (relative to incident irradiance) and O_2_ concentration on Day 4.

On Day 4, the 2D cross-section of the simulated light field in the construct (Figure 5d, figure S11a) and the corresponding 1D line profiles (Figure 5f, Figure S11c) showed that in slab_PS, light was almost fully attenuated within the top 500 µm. In contrast, light penetrated much deeper in the PS reaching nearly the entire depth near the water channels (Figure 5d), though light was still strongly attenuated within the top 500 µm in regions farther from the channels (Figure S5d and Figure S11a). Despite these differences in light penetration, the average (simulated) scalar irradiance for PS on Day 4 was similar to that of slab_PS (∼96%, bar #5 in Figure S6a).

The 2D cross-section showing the simulated steady-state O₂ distribution in the constructs for Day 4 (Figure 5e, Figure S11b) and the corresponding 1D depth profiles extracted along a line (Figure 5f, Figure S11c) showed a slightly higher O₂ accumulation in the deeper regions of PS compared to slab_PS. This is likely due to the deeper light penetration in PS and its higher net O₂ production rate (Section 3.3 and Table T2).

Although PS exhibited a 2.2-fold higher net O₂ production rate (Figures S6a, 4a, 4b), its average steady-state O₂ concentration (simulated) was only marginally higher than that of slab_PS (PS = 105% of slab_PS) in the simulations, due to its more efficient mass transfer with the surrounding medium. The significantly higher algal growth rate observed in bioprinted PS constructs (2.4 times that of slab_PS) was thus likely the result of this combination of improved mass transfer and a larger illuminated surface area.

#### 3.4.2 Vgroove versus slab_Vgroove

On Day 0, the 2D cross-section of the simulated light field (Figure 6a) and the corresponding 1D profiles extracted along a vertical line (Figure 6c, S12) showed that light penetrated throughout the entire Vgroove sample. The average scalar iiradiance (simulated) for Vgroove was similar to that of slab_Vgroove (94% of the slab value, bar #5 in Figure S6b). However, the Vgroove construct had twice the illuminated surface area as compared to the corresponding slab_Vgroove. The 2D cross-section of the simulated steady-state O₂ distribution (Figure 6b) and the corresponding 1D profiles (Figure 6c, S12) revealed a higher O₂ accumulation in the slab_Vgroove constructs as compared to the Vgroove construct. This difference is likely due to a more efficient mass transfer between the Vgroove construct and the surrounding medium, which can be attributed to its surface area-to-volume ratio being twice that of the slab. As a result, the overall average steady-state O₂ concentration (simulated) in Vgroove on Day 0 was approximately 70% of that in slab_Vgroove (Figure S6b).

**Figure 6:**
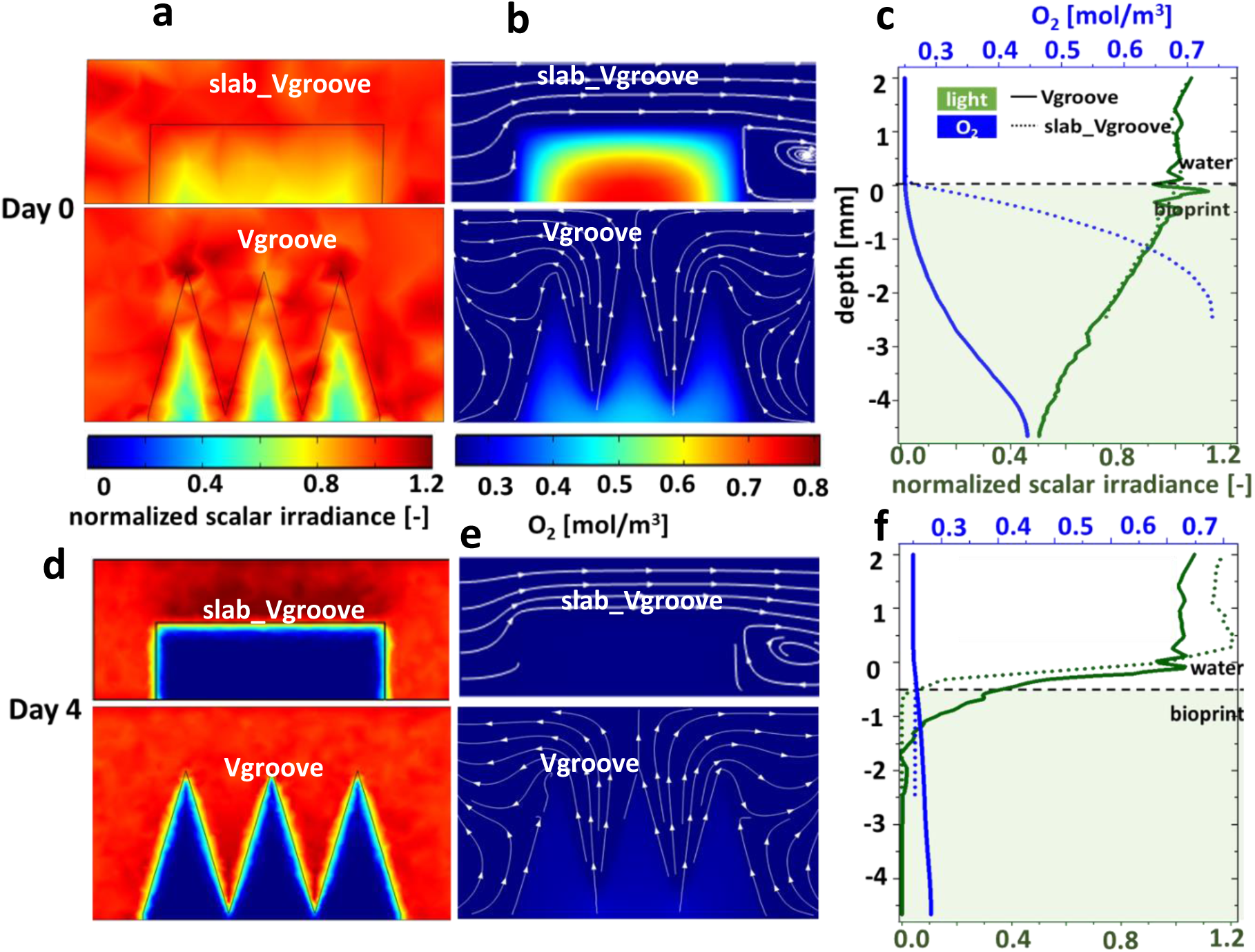
Simulation of steady state light and O_2_ distribution in bioprinted Vgroove and slab_Vgroove constructs. **a:** 2D cross-section (see figure S8) showing the simulated vertical light distribution (normalized scalar irradiance 636 nm) on Day 0; **b:** 2D cross-section of the corresponding O_2_ distribution on Day 0; **c:** Extracted line profiles (see figure S8) of normalized scalar irradiance (relative to incident irradiance) and O_2_ concentration on Day 0. **d:** 2D cross-section (see figure S8) showing the simulated vertical light distribution (normalized scalar irradiance for 636 nm) on Day 4; **e:** 2D cross-section of the corresponding O_2_ distribution on Day 4. **f:** Extracted line profiles (see figure S8) of normalized scalar irradiance (relative to incident irradiance) and O_2_ concentration on Day 4.

On Day 4, the 2D cross section of the simulated light field (Figure 6d) and the corresponding 1D profiles (Figure 6f, Figure S12) showed that light was almost completely attenuated within the top 500 µm of slab_Vgroove. In contrast, light penetration was deeper in the Vgroove, reaching approximately 1.5 mm near the peaks (Figure 6f) and about 1 mm from the slanted edges (Figure S12), where increased reflective losses occurred at the angled surfaces. As a result, the average (simulated) scalar irradiance in Vgroove was 37% higher than in the slab (Figure S6b). The 2D cross-section showing simulated steady-state O₂ distribution (Figure 6e) and the corresponding 1D profiles (Figure 6f, S12) showed a higher O₂ accumulation in deeper regions of the Vgroove construct, as compared to slab_Vgroove. This was driven by by the higher light availability causing higher net O₂ production rates. Despite Vgroove exhibiting a 10-fold increase in net O₂ production (Figures S6b, 4c, 4d), the average steady-state O₂ concentration (simulated) remained comparable to that of slab_Vgroove (Vgroove = 106% of slab_Vgroove), due to more efficient mass transfer. The 1.3-fold higher algal growth rate observed in Vgroove in constructs was thus likely the result of a combination of factors: improved mass transfer, increased illuminated surface area, and higher overall light availability as biomass increased (w.r.t. slab_Vgroove on Day 4), compared to the slab.

Under the conditions given during the first 4 days of incubation, the bioprinted PS construct exhibited a higher algal growth rate, i.e., 2.4 times that of slab_PS, while the bioprinted Vgroove construct exhibited 1.3-fold algal growth rate than in the bioprinted slab_Vgroove. This difference can be attributed to the PS construct having a higher surface area-to-volume ratio (2.5 times that of the slab) than the Vgroove construct (2 times the slab). The greater surface area in the PS construct enhanced both light exposure and mass transfer, not only for O₂, but also for other key species such as CO₂ and nutrients, which collectively contribute to increased algal growth rates.

In addition to influencing light distribution, we note that the shape of the bioprinted construct can alter flow patterns around the sample, thereby affecting mass transfer. As the simulations in this study are intended as a guiding tool to help interpret experimental observations, they do not precisely replicate the hydrodynamic conditions during incubation or gas exchange measurements. While the experiments were conducted under stirred conditions, the simulations assume a simplified laminar flow scenario. Future work will focus on modeling more complex flow dynamics and integrating them with photosynthetic activity.

## 4. Conclusions and Outlook

We present a proof-of-concept framework that integrates predictive modeling, experimental validation, and simulation to evaluate and optimize the performance of microalgal bioprints. By incorporating experimentally measured photosynthetic parameters into simulations, we were able to gain mechanistic insights into how structural design influences algal growth dynamics. Specifically, we demonstrate that bioprinted construct geometries such as PS and Vgroove structures significantly enhance algal growth rates, as compared to flat bioprinted slab geometries representative of simple biofilms. This improvement could be attributed to increased light penetration, a higher surface area-to-volume ratio, and enhanced mass transfer with the surrounding medium. Our study paves the way for the rational design of more efficient bioprinted algal construct geometries for use in surface-associated bioproduction (e.g. in photobioreactors). Such structured geometries hold immense potential for maximizing algal productivity in compact formats, addressing critical limitations of conventional systems that require large land areas^2,4^. The ultimate goal is to engineer space-efficient bioprint designs that optimize light capture and nutrient exchange, enabling high-density algal cultivation within a minimal land footprint, an essential step toward scalable, sustainable biofuel and bioproduct generation.

The demonstrated fabrication, measurement and simulation pipeline can be used to study other structure-function relationships seen in phototrophic organisms like corals^14,22,55,56^, jelly fish^57^ and other photosymbiotic organisms. The capacity to replicate complex light–mass transfer interactions in such systems could shed light on ecological adaptations and energy optimization strategies in nature. Furthermore, this platform holds promise for applications in synthetic biology and bio-inspired design. By integrating multiple cell types and layered structures, it can be used to engineer bionic constructs to study symbiotic relationships^33^, microbe-host interactions^34–36^, and metabolic co-dependencies. These capabilities may also inform therapeutic design, targeted drug delivery, and tissue engineering applications. Future work will focus on expanding this platform to include multilayered 3D bioprints e.g. inspired by the hierarchical organization of reef-building corals^58^ or complementary spectral absorption in stratified microbial biofilms^59,60^, enabling spatial separation and canopy formations of different cell types with complementary functions. Additionally, we aim to incorporate more realistic hydrodynamic conditions, including turbulent and oscillatory flows commonly found in natural environments, to better replicate *in vivo* performance. Incorporating adaptive or responsive materials could further allow dynamic modulation of mass and light transfer, ultimately enabling the development of highly optimized, self-regulating bioreactors.

## Author Contributions

SM and MK conceptualized and designed the research. SM and MM synthesized GelMA. SM prepared the bionk, optimized printing and photocuring, and fabricated the bioprints. SM performed OCT imaging and analysis, cell counts in bioprinted constructs, respirometry measurements and image analysis. SM performed numerical simulations. SM and MK wrote the manuscript with editorial input from MM.

## Acknowledgements

We thank Sofie Lindegaard Jakobsen and Veronica Bach Petersen for excellent technical assistance. The study was supported by research grants from the Villum Foundation [VIL50371 & VIL57413; MK], the Independent Research Fund Denmark [DFF-8022-00301B; MK], the European Union (Marie Skłodowska-Curie Grant Agreement No. 101073507; MK), and the Gordon and Betty Moore Foundation [grant no. GBMF9206; https://doi.org/10.37807/GBMF9206; MK].

## Supplementary information

**Table T1:**
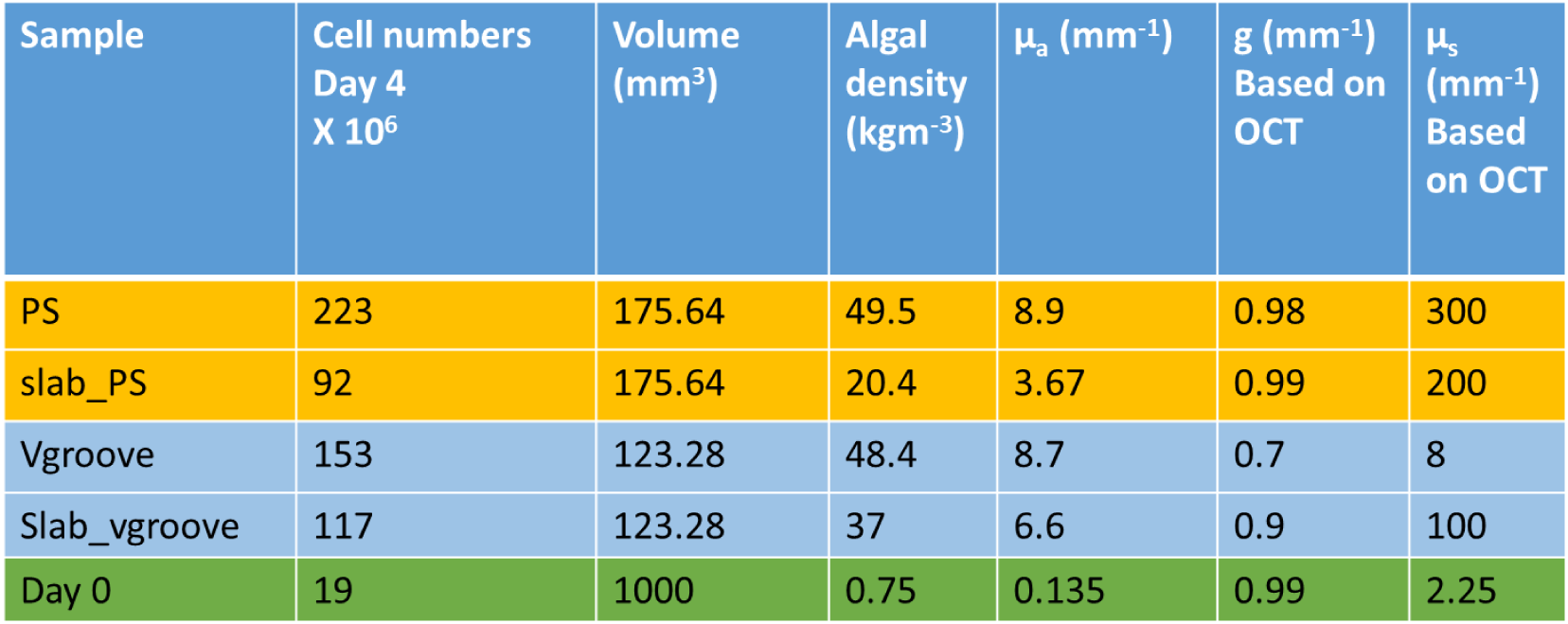
Estimated optical properties of the samples on Day 4, as determined from the OCT images shown in Figure 3c and 3d and cell counts. The lst row in the table lists the optical properties on Day 0, as estimated from the cell count.

**Table T2:** Model parameters for flow, mass transport and biokinetics.

| Parameter | Symbol | Value | Unit | Source |
| --- | --- | --- | --- | --- |
| **Flow** |  |  |  |  |
| Viscosity water | $\eta_{wat}$ | 0.0009 | Pa s | ref<sup>61</sup> |
| **Densities** |  |  |  |  |
| water | $\rho_{wat}$ | 997 | kg m<sup>-3</sup> | ref<sup>61</sup> |
| bio-print | $\rho_{bp}$ | 1109 | kg m<sup>-3</sup> | ref<sup>62</sup> |
| **Mass transport** |  |  |  |  |
| **Diffusion coefficients** |  |  |  |  |
| O<sub>2</sub> in water | $D_{O_2,wat}$ | $2 \cdot 10^{-9}$ | m<sup>2</sup> s<sup>-1</sup> | ref<sup>63</sup> |
| O<sub>2</sub> in bio-print | $D_{O_2,bp}$ | $1 \cdot 10^{-9}$ | m<sup>2</sup> s<sup>-1</sup> | ref<sup>64</sup> |
| **Experimental parameters** |  |  |  |  |
| Average water velocity | $v_0$ | 5 | mm s<sup>-1</sup> | ref<sup>14</sup> |
| Ambient temperature | $T_0$ | 25 | °C | ref<sup>14</sup> |
| Inlet O<sub>2</sub> concentration | $c_{0,O_2}$ | 0.25 | mol m<sup>-3</sup> | ref<sup>65</sup> |
| Incident downwelling photon irradiance (400 – 700 nm) | $I_0$ | 430 | μmol photons m<sup>-2</sup> s<sup>-1</sup> | |
| Net-quantum efficiency_Day0; all samples | $net-QE_{day0}$ | $5 \cdot 10^{-3}$ | - | |
| Net-quantum efficiency_Day4_PS | $net-QE_{PS}$ | $5 \cdot 10^{-4}$ | - | |
| Net-quantum efficiency_Day4_slab_PS | $net-QE_{slab\_PS}$ | $5.4 \cdot 10^{-4}$ | - | |
| Net-quantum efficiency_Day4_Vgroove | $net-QE_{Vgroove}$ | $2.2 \cdot 10^{-4}$ | - | |
| Net-quantum efficiency_Day4_slab_Vgroove | $net-QE_{slab\_Vgroove}$ | $4.4 \cdot 10^{-5}$ | - | |

**Figure S1:**
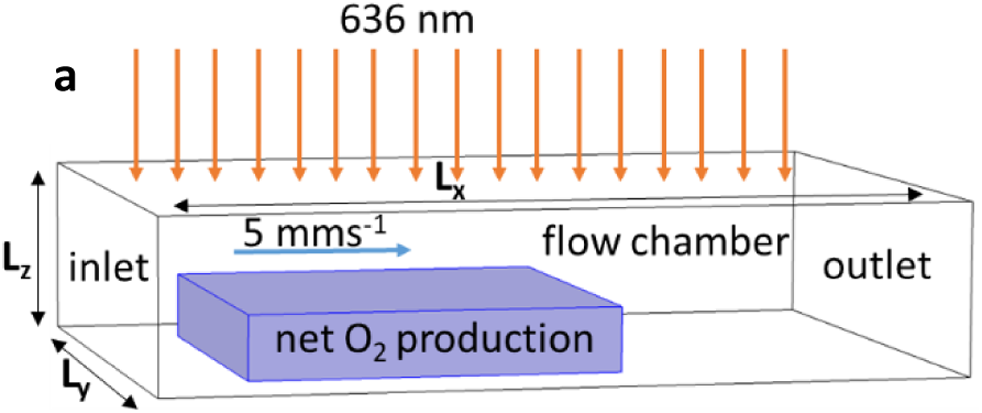
Sketch showing the model geometry of slab in a flow chamber used for light, flow and mass transfer simulation. Downwelling irradiance at 636 nm illuminates the entire top boundary of the flow chamber. No light flux condition to all other boundaries. The dimensions of the bioprints are as shown in Figure 1. Simulations assumed a fully developed laminar flow with an average velocity of 5 mm s^−1^ and a constant O_2_ concentration at the inlet. A fixed gauge pressure of zero and convection only condition was assumed at the outlet. We assumed a o slip boundary condition at the water bio-print interface, and symmetry and ‘no flux’ boundary conditions on the top and lateral walls of the flow chamber. A net photosynthetic oxygen production rate, calculated from the light field (MC simulation) within the biofilm, was assigned to the bioprint. L_x_ bioprint length + 12 mm, L_y_ bioprint width + 4mm, L_z_ bioprint height + 2 mm.

**Figure S2:**
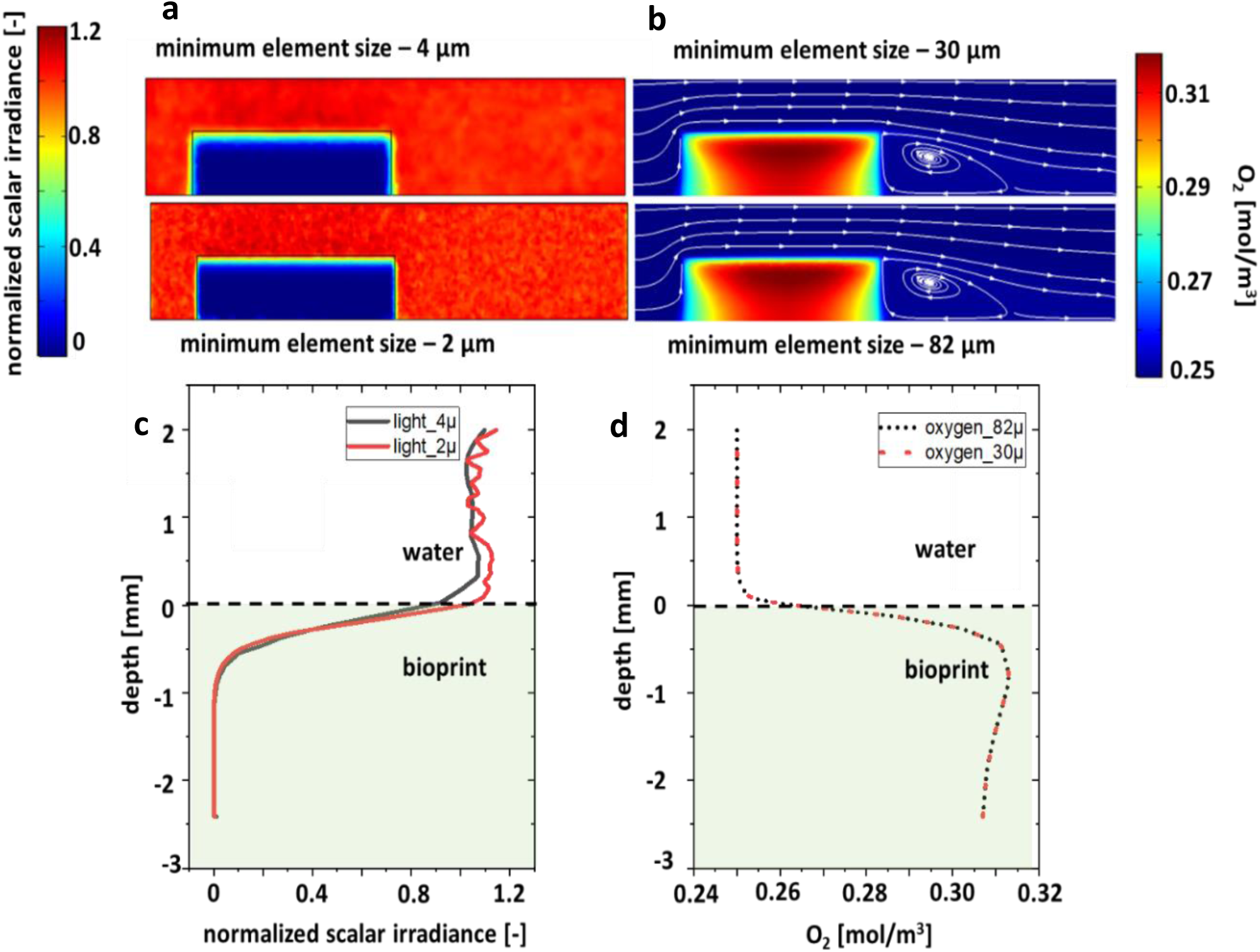
Mesh independent study for slab_PS on day 4. **a:** 2D cross-sectional and **c:** 1D plot of normalized scalar irradiance showing that the simulated light field (at 636 nm) is similar for meshes with minimum element size of 2 µm and 4 µm, respectively. A 4 µm minimum element size mesh was used for light simulations for the entire study. **b:** 2D cross-sectional and **d:** 1D plot showing that the simulated O_2_ concentration (for downwelling irradiance of 430 μmol photons m^−2^ s^−1^ integrated over 400 – 700 nm) is identical for minimum element size of 30 µm and 82 µm. A 82µm minimum element size mesh was used for O_2_ mass transfer simulation for the entire study.

**Figure S3:**
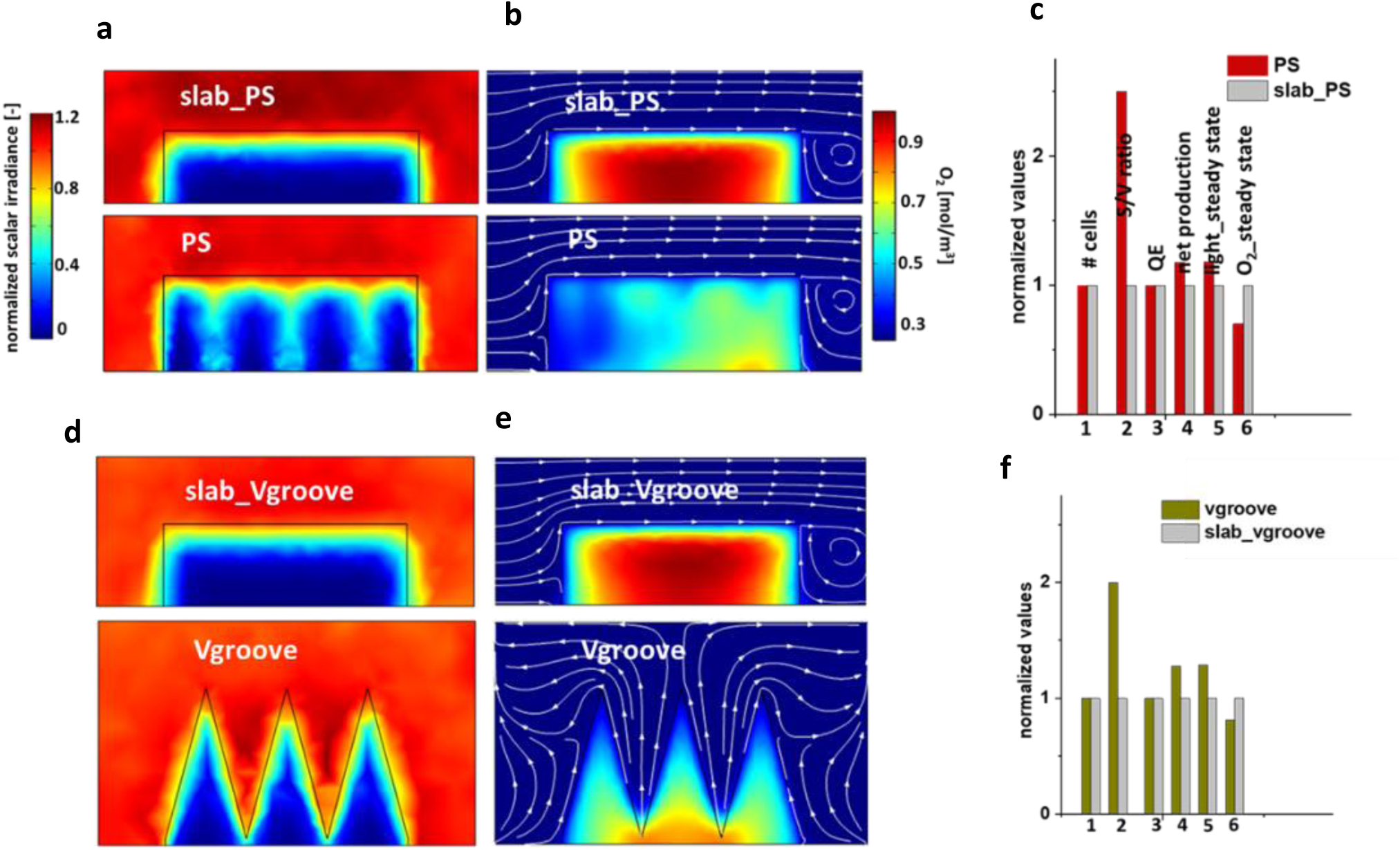
Steady state light and O_2_ mass transfer simulations with assumed optical properties. (µ_a_ 1.7 mm^−1^, µ_s_ 90 mm^−1^, g 0.98, n 1.33 at 636 nm) and photosynthetic net quantum efficiency 0.005 at an incident photon irradiance of 320 µmol photons m^−2^ s^−1^ (400-700 nm). **a:** 2D cut plane data (Figure S7) of light simulation of PS and slab_PS; **b:** 2D cut plane data of corresponding O_2_ mass transfer simulation for PS, slab_PS; **c:** Comparison between PS and slab_PS showing 17% higher light availability and 30% lesser oxygen accumulation for PS w.r.t slab_PS; **d:** 2D cut plane data (figure S8)of light simulation of Vgroove and slab_Vgroove; **e:** 2D cut plane data of corresponding O_2_ mass transfer simulation for Vgroove, slab_Vgroove; **f:** Comparison between Vgroove and slab_Vgroove showing 27% higher light availability and 20% lesser oxygen accumulation for Vgroove w.r.t slab_Vgroove.

**Figure S4:**
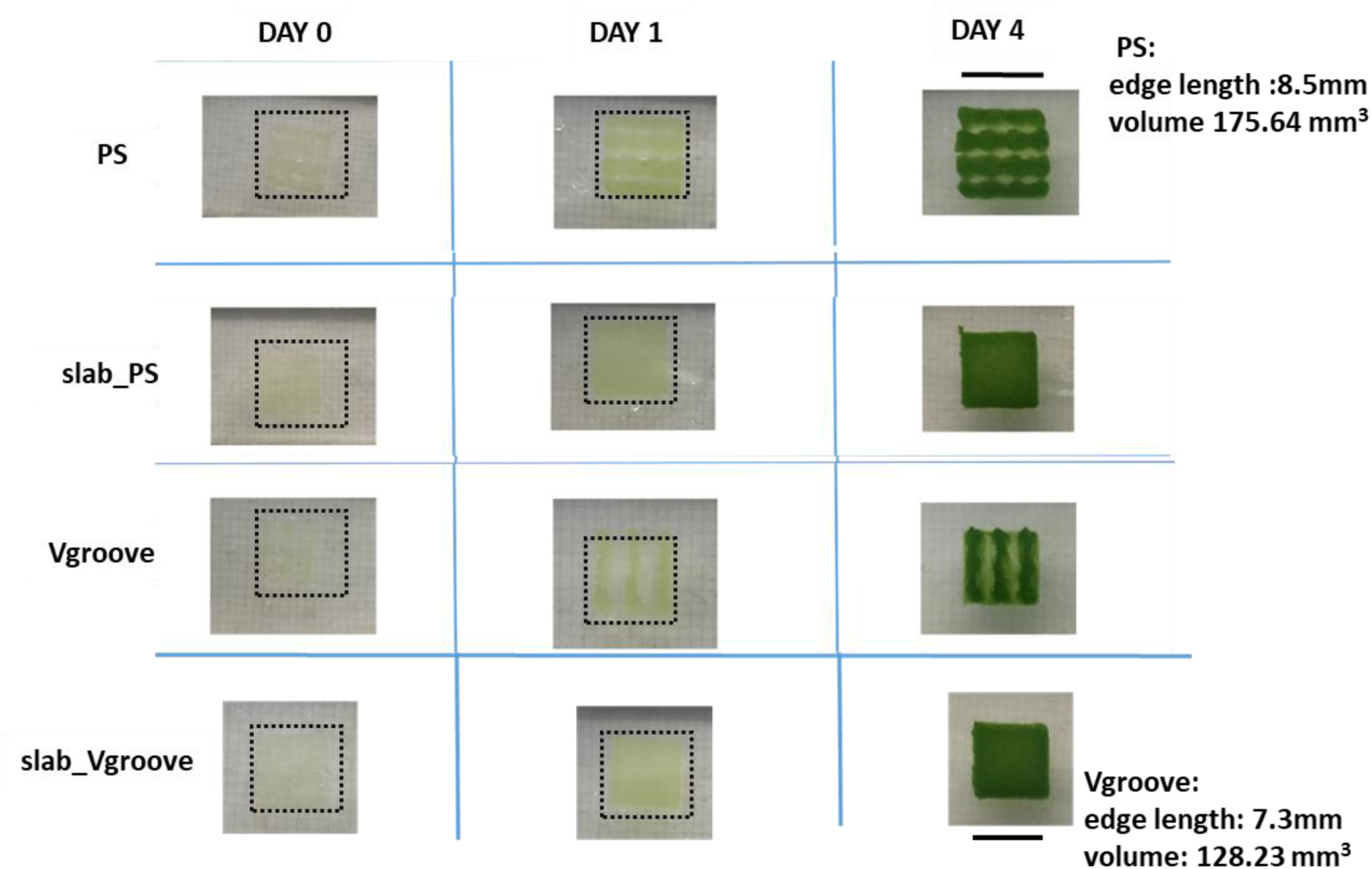
Photographs of the samples, stored in TAP medium, taken on Day 0, Day 1 and Day 4.

**Figure S5:**
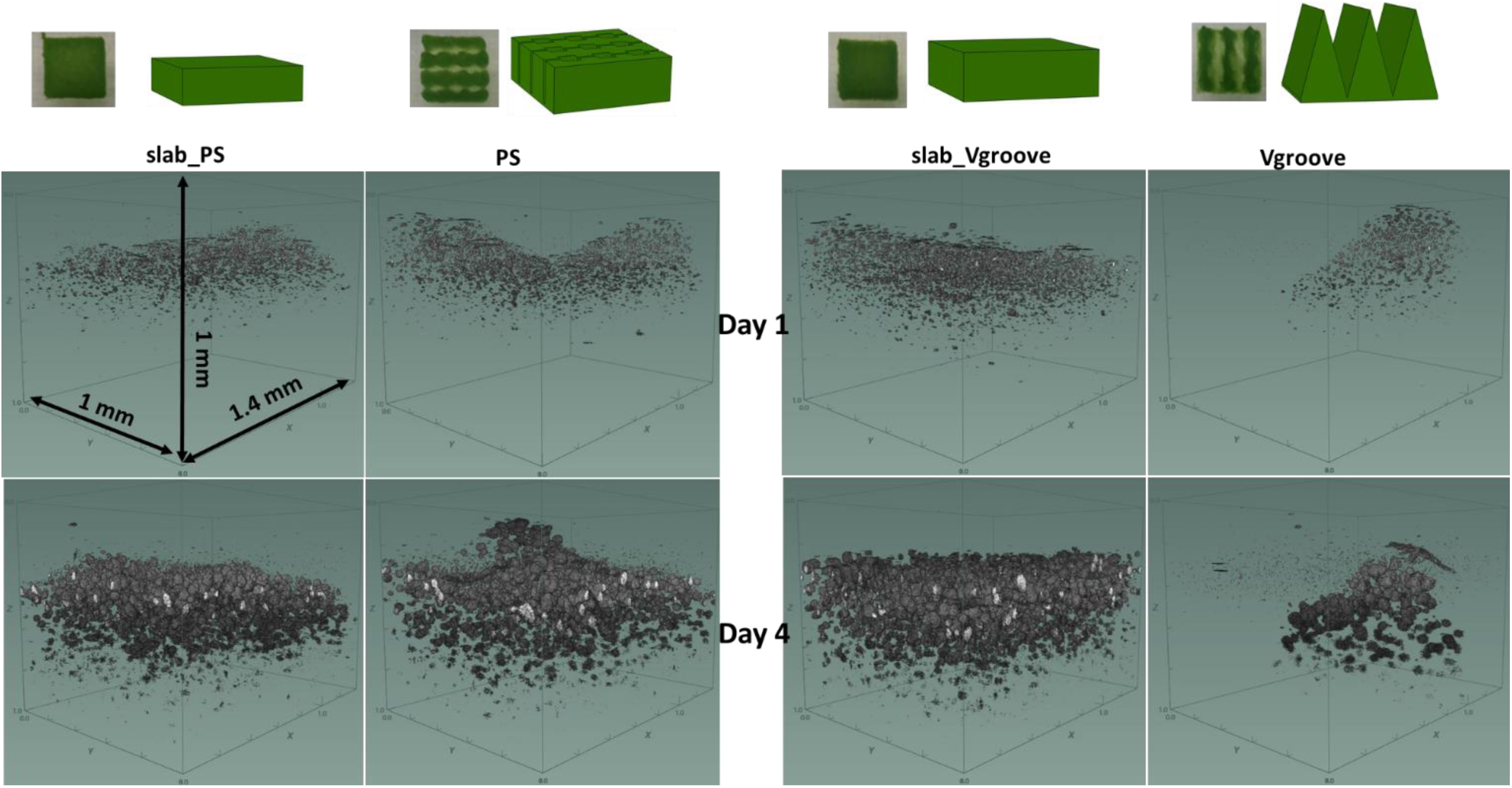
3D rendering of optical coherence tomography (OCT) images of the different bioprinted geometries, shown in figure 4, on Day 1 and Day 4. See also the videos of slab_PS from Day 1 and Day 4, which show the aggregation of algal cells.

**Figure S6:**
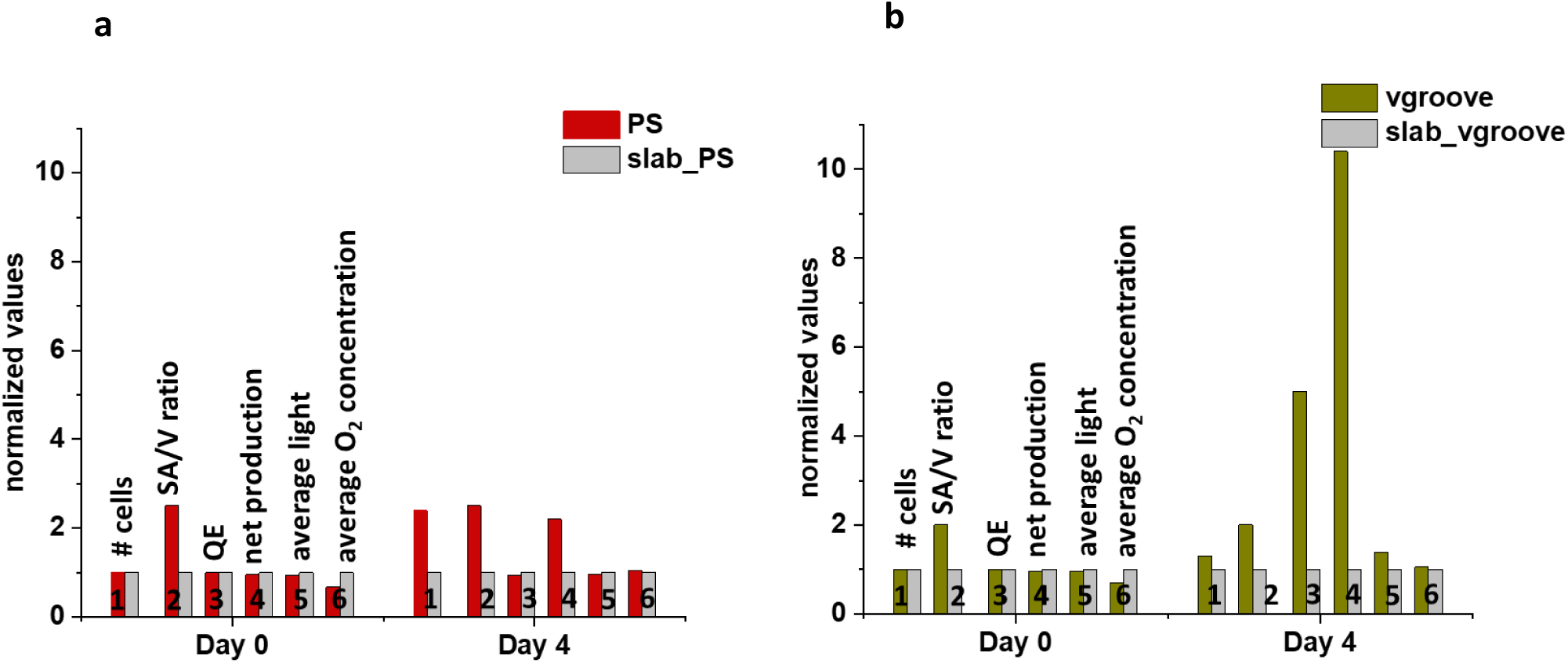
Comparison of various simulated parameters between the different bioprinted constructs and their respective slabs on Day 1 and Day 4, for. **a:** PS and slab_PS; **b:** Vgroove and slab_Vgroove. The light and O₂ values shown represent averages across the entire sample and are normalized to the corresponding slab, highlighting the differences between each 3D design and its slab counterpart.

**Figure S7:**
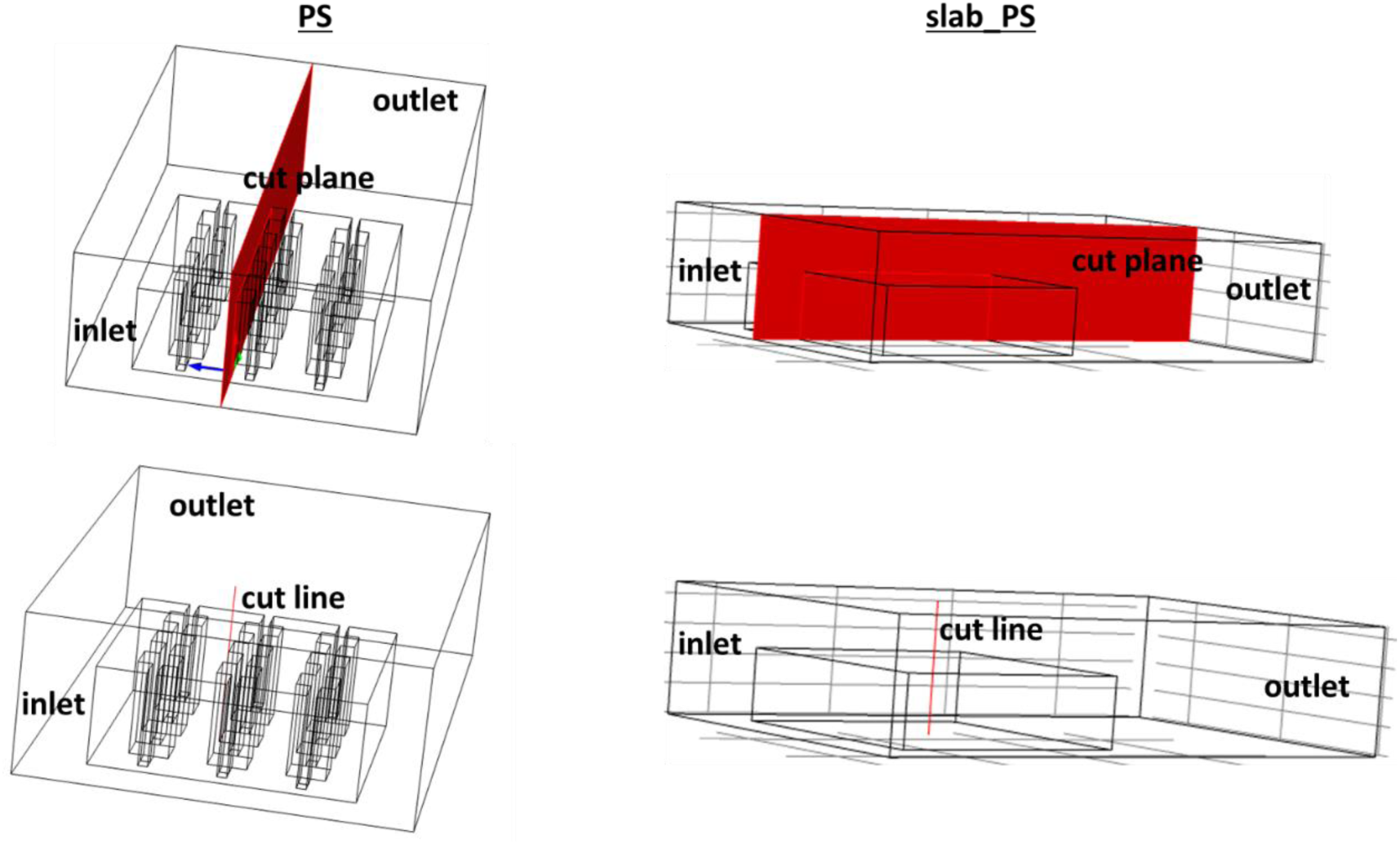
Schematic showing the position of cut plane and line for extraction of the simulation data shown in figure 5 for PS and slab_PS.

**Figure S8:**
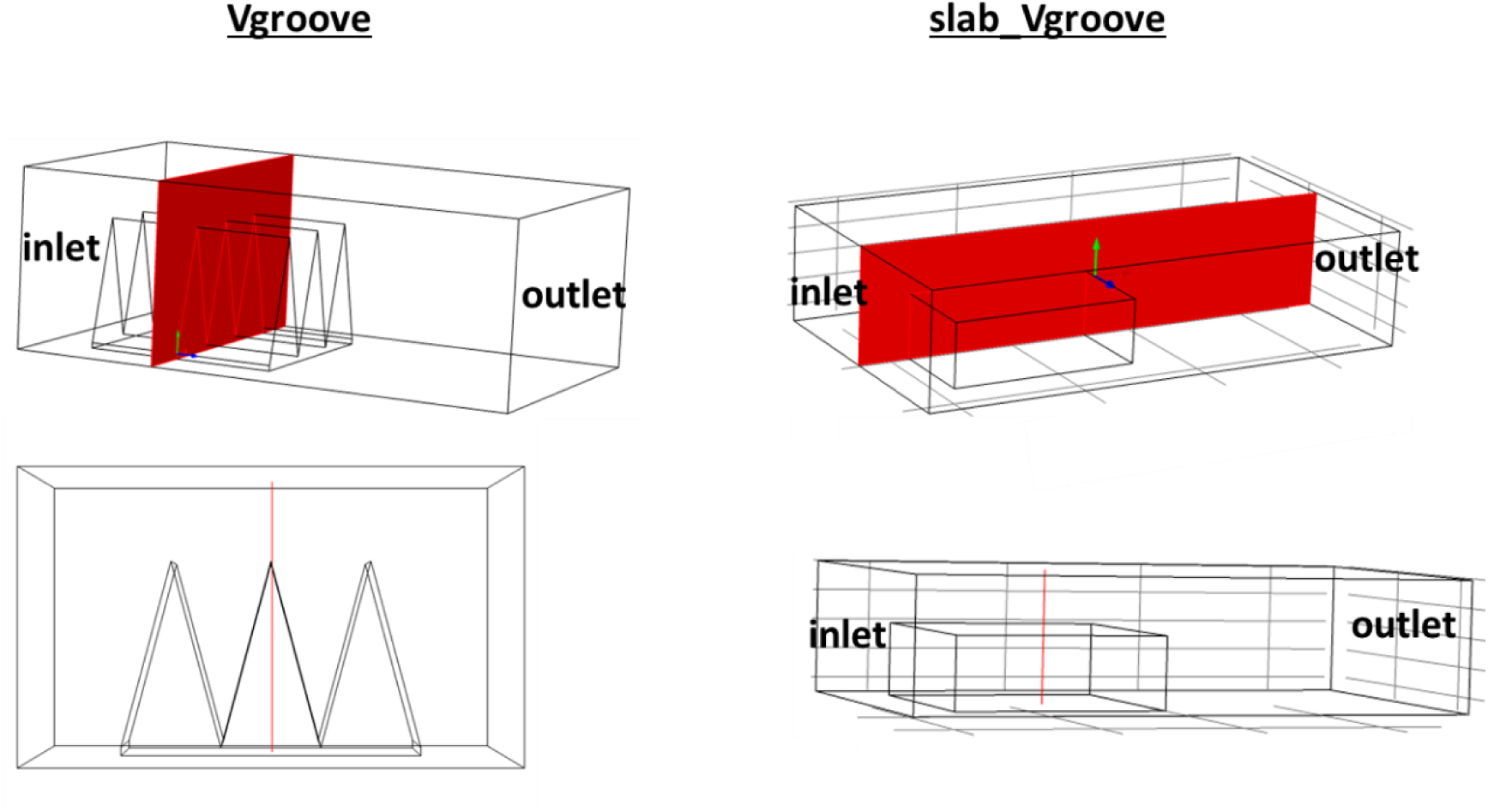
Schematic showing the position of cut plane and cut line for extraction of the simulation data shown in figure 6 for Vgroove and slab_Vgroove.

**Figure S9:**
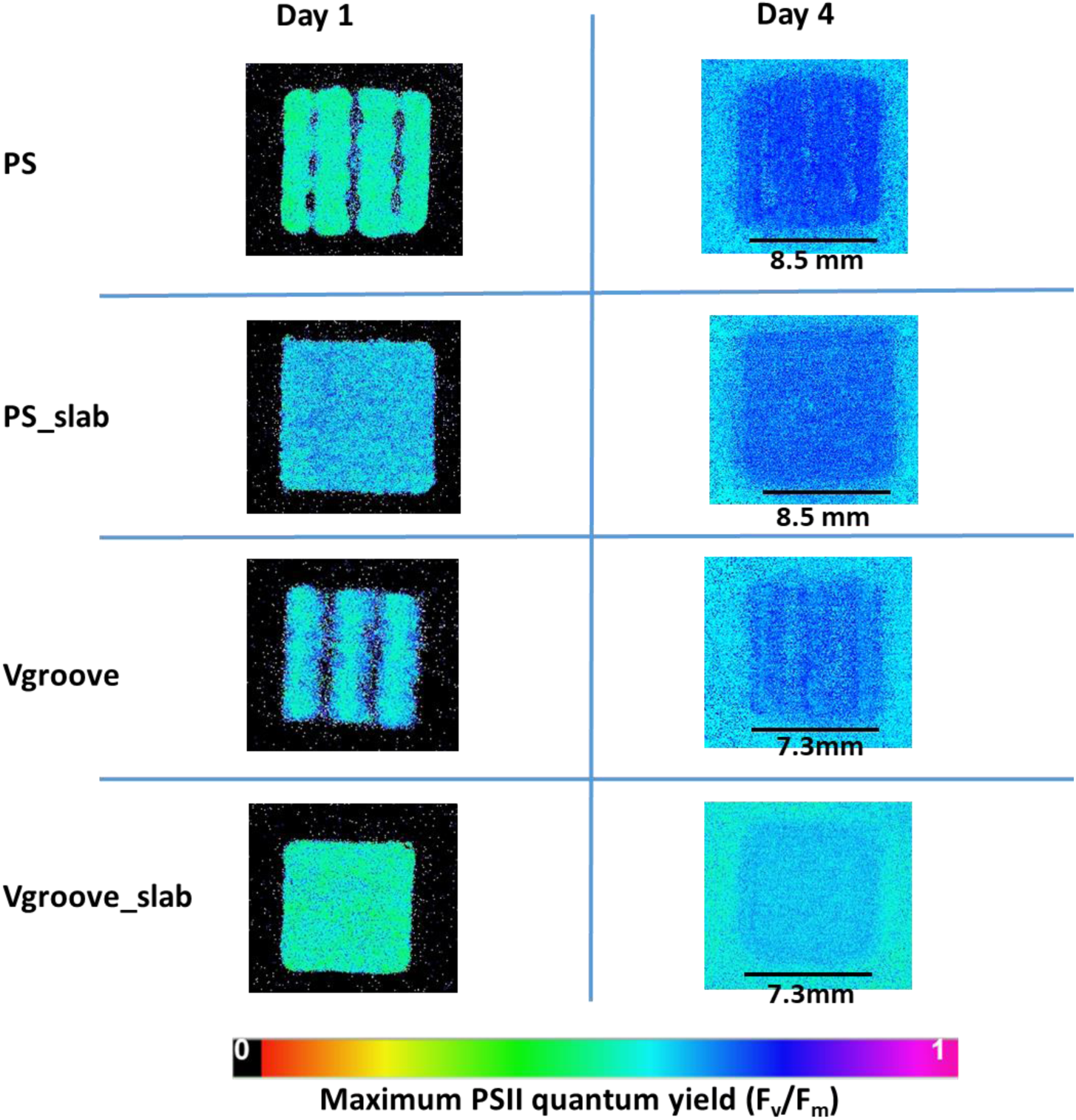
Maximum PSII quantum yield (F_v_/F_m_) images of all the samples on Day 1 and Day 4.

**Figure S10:**
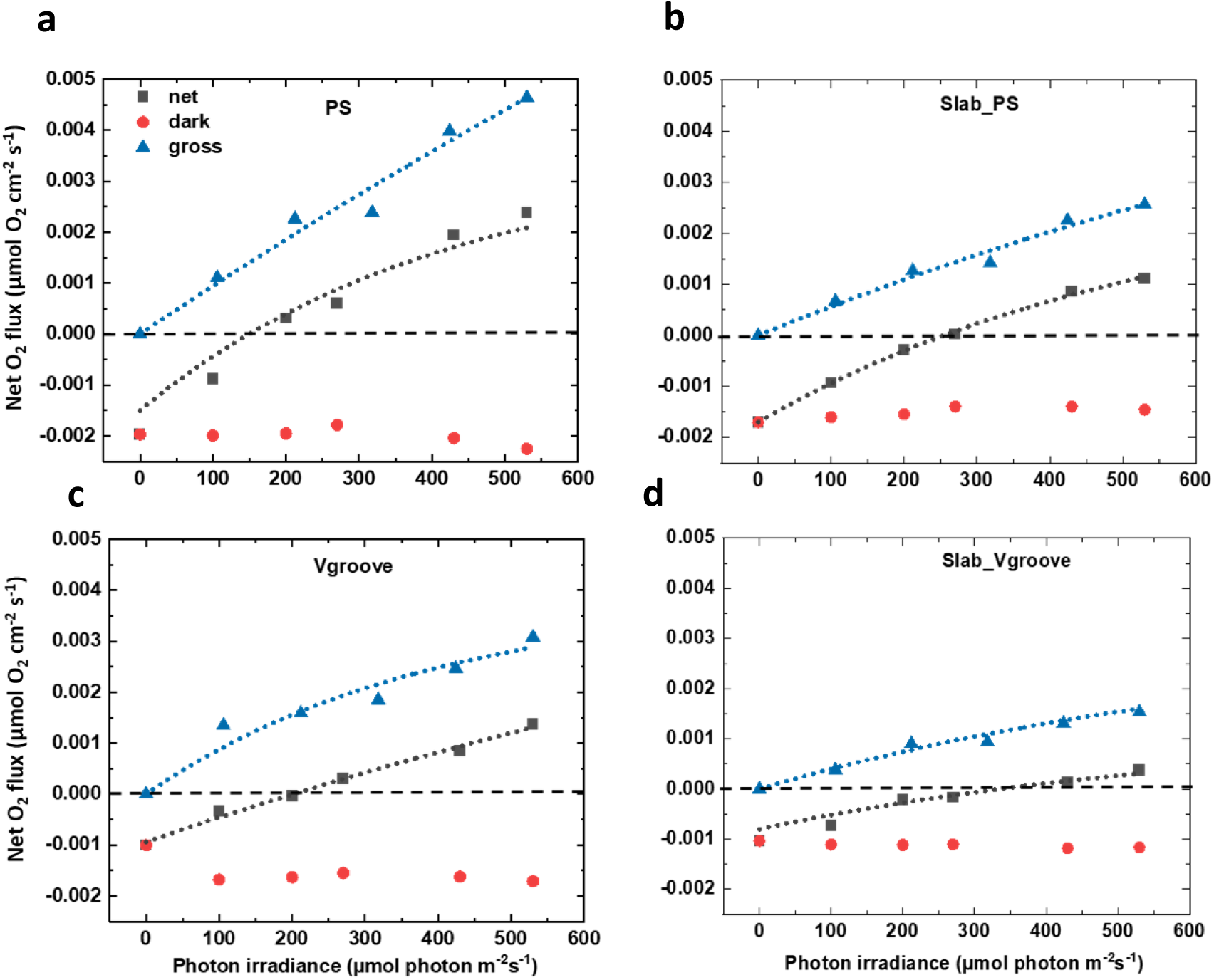
Net and estimated gross photosynthesis rates and dark respiration rates for different bioprinted construct geometries. **a:** PS; **b:** Slab_PS, **c:** Vgroove; **d:** Slab_Vgroove; as determined from the slopes in Figure 4a and 4c (symbols). The net production as a function of photon irradiance (400 – 700 nm) was fitted according to Spilling et al.^1^ and the estimated gross production according to Webb et al.^5^(dotted lines). The rates are normalized to construct footprint area.

**Figure S11:**
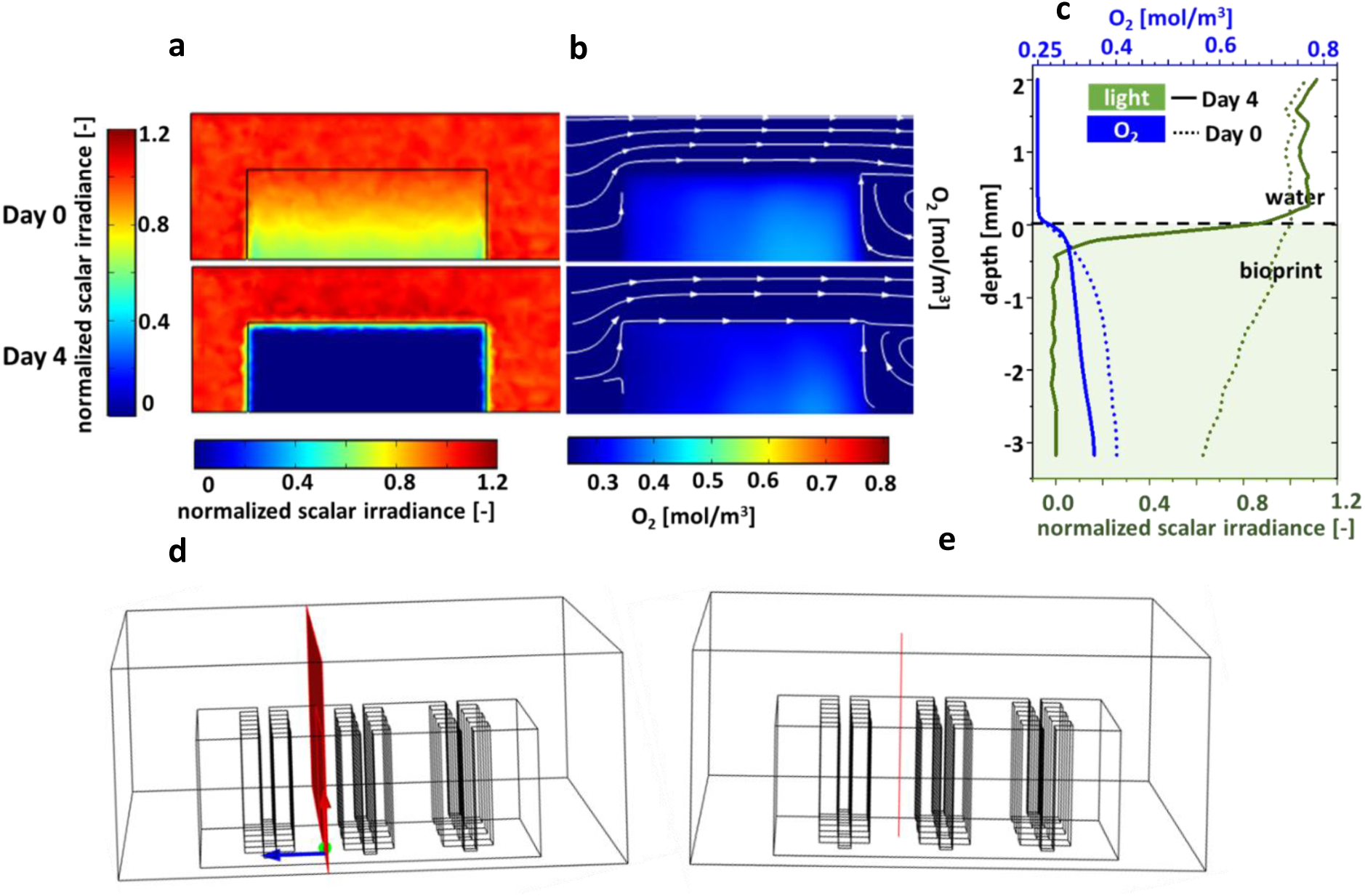
Steady state light and O_2_ mass transfer simulation results for PS and slab_PS constructs along the middle plane. **a:** 2D cut plane data of light simulation on Day 0 and Day 4; **b:** 2D cut plane data of corresponding O_2_ mass transfer simulation on Day 0 and Day 4; **c:** Cut line data of both light and O_2_ mass transfer simulations on Day 0 and Day 4. **d:** Schematic showing position of 2D cut plane; **e:** Schematic showing the position of cut line.

**Figure S12:**
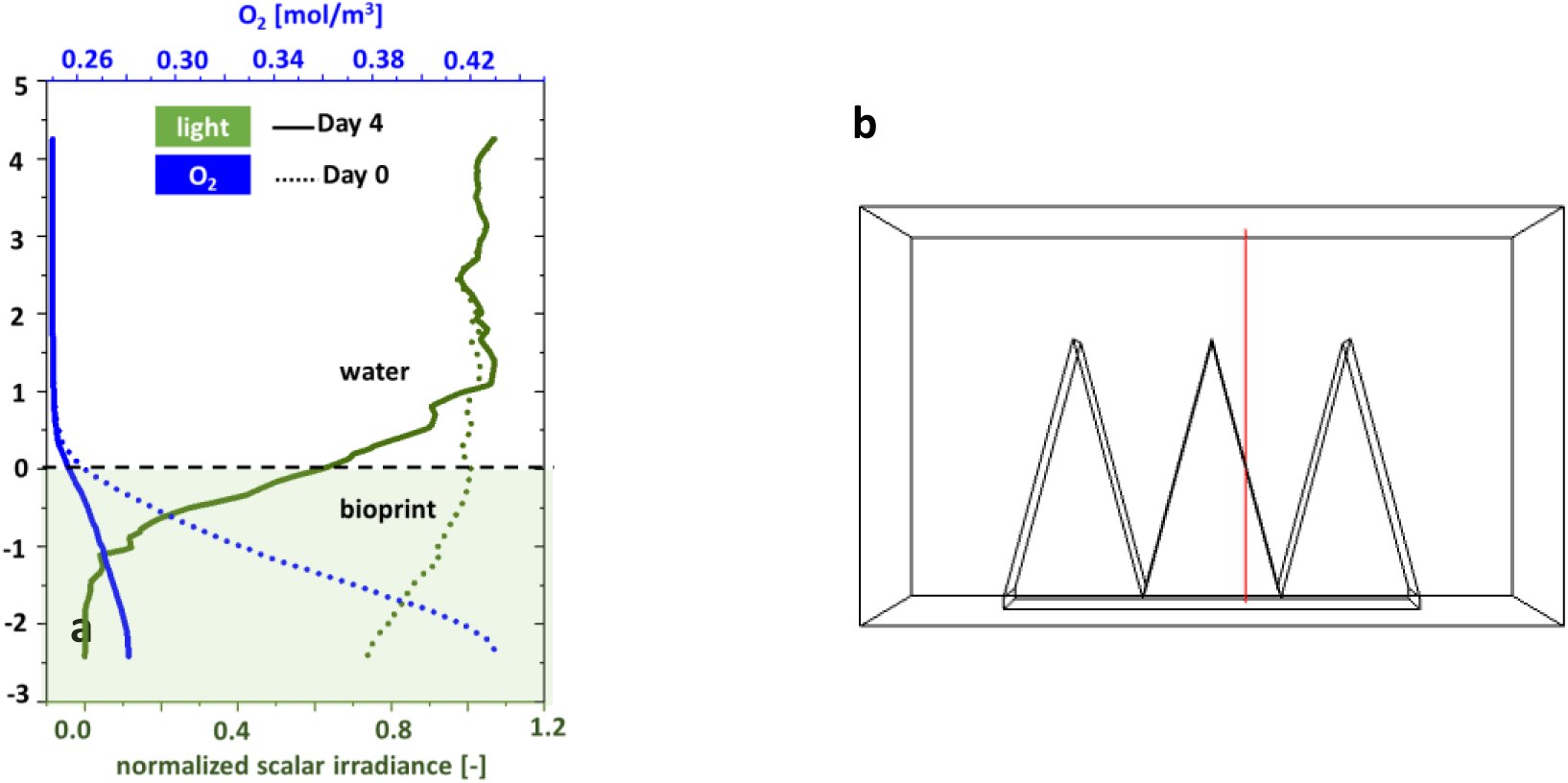
Steady state light and O_2_ mass transfer simulation results for Vgroove. **a:** Cut line data of both light and O_2_ mass transfer simulations on Day 0 and Day 4. **b:** Schematic showing the position of the cut line.

